# Cross-ploidy hybridisation in Alpine woodrushes is associated with ecological additivity and scale-dependent niche divergence

**DOI:** 10.64898/2026.02.26.708247

**Authors:** Valentin Heimer, Peter Schönswetter, Božo Frajman

**Affiliations:** Department of Botany, University of Innsbruck, Sternwartestraße 15, 6020 Innsbruck, Austria; Institute for Alpine Environment, Eurac Research, Drususallee 1/Viale Druso 1, 39100, Bozen/Bolzano, Italy

**Keywords:** cross-ploidy hybridisation, ddRADseq, Eastern Alps, ecological additivity, *Luzula*, niche evolution, polyploidy, morphometry

## Abstract

- Hybridisation and whole-genome duplication (WGD) are widespread in plants, yet their ecological consequences remain challenging to predict. In allopolyploids, where both processes coincide, ecological divergence is typically evaluated against a null hypothesis of ecological additivity. However, this hypothesis has not been tested explicitly in the context of cross-ploidy hybridisation.
- We assessed ecological additivity following cross-ploidy hybridisation in the Alpine woodrush *Luzula alpina* across multiple spatial resolutions. Integrating genomic data, environmental niche modelling, vegetation relevés, and morphometric analyses, we characterised population structure and quantified ecological and morphological differentiation between the hybrid and its parental species.
- Genomic and morphometric evidence confirmed the hybrid origin of *L. alpina*. Patterns of niche evolution varied with spatial resolution. Climatic and edaphic data supported ecological additivity, whereas plot-level vegetation data revealed subtle but significant niche divergence accompanied by shifts in associated plant community composition. Postglacial population histories further suggest long-term persistence of hybrid and parental lineages in distinct refugia followed by differential recolonisation of the Alps.
- Our results indicate that cross-ploidy hybridisation in Alpine *Luzula* is predominantly associated with niche stability rather than pronounced ecological divergence. Importantly, the detection of divergence beyond ecological additivity depends on the spatial resolution of environmental data.

## Introduction

Hybridisation is a major driver of speciation across plants and animals (Rieseberg & Willis, 2007; Soltis & Soltis, 2009; Abbott *et al*., 2013), with at least 25% of plant species estimated to participate in hybridisation (Mallet, 2005). Homoploid hybrid speciation (HHS) occurs without changes in chromosome number and only a few unequivocal cases of HHS have been documented (Schumer *et al*., 2014; Yakimowski & Rieseberg, 2014; Nieto Feliner *et al*., 2017). On the other hand, allopolyploid speciation that involves hybridisation coupled with whole-genome duplication (WGD) is widely acknowledged as a primary mechanism of plant diversification (Soltis & Soltis, 2009; Wood *et al*., 2009).

Irrespective of their mode of origin, hybrid species must achieve reproductive isolation from their progenitors to become established. In allopolyploids, WGD generates strong postzygotic barriers, conferring immediate reproductive isolation (Levin, 2002; Ramsey & Schemske, 2002). In contrast, homoploid hybrids lack such inherent reproductive isolation and must rely on alternative mechanisms to reduce gene flow. These include prezygotic barriers such as shifts in mating system, flowering phenology, or pollination syndrome (Lowe & Abbott, 2004; Melo *et al*., 2009; Runemark *et al*., 2019), as well as postzygotic barriers arising from chromosomal rearrangements (Rieseberg *et al*., 1995; Lai *et al*., 2005). The phenotypic consequences of hybridisation are not restricted to reproductive barriers but more generally result in morphological intermediacy (Wilson, 1992; Čertner *et al*., 2015), or mosaic combinations of parental and intermediate traits (Rieseberg *et al*., 1993; Skubic *et al*., 2023).

Beyond intrinsic barriers, reproductive isolation may be reinforced by ecological divergence, whereby hybrid lineages exploit novel niches (Rieseberg *et al*., 2003; Gompert *et al*., 2006). Ecological differentiation is also crucial for the establishment of neoallopolyploids, which escape minority cytotype exclusion through either geographical or ecological isolation from their progenitors (Levin, 1975, 2003; Fowler & Levin, 1984). Although competitive exclusion of parental species within conserved niches is theoretically possible, it is rarely observed as parental species are already established and adapted to the local conditions.

Consequently, niche shifts in allopolyploids are considered the rule rather than the exception (Blaine Marchant *et al*., 2016; Baniaga *et al*., 2020; Akiyama *et al*., 2021; Lu *et al*., 2025). Polyploids have long been hypothesised to be superior colonisers of novel or harsh environments (Fowler & Levin, 1984; Leitch & Leitch, 2008; Shimizu-Inatsugi *et al*., 2017), yet empirical data do not show conclusive trends. Despite the higher frequency of polyploids at high latitudes (Rice *et al*., 2019), recent analyses have found no consistent climatic niche shifts in autopolyploids relative to their diploid progenitors (Meirmans & Kolář, 2025), underlining the uncertainty about the ecological consequences of WGD.

In allopolyploids, disentangling the ecological effects of WGD from those of hybridisation is particularly challenging (Meirmans & Kolář, 2025). Because of their fixed heterozygosity, allopolyploids are expected to occupy the combined niche space of their progenitors, a prediction formalised as the null hypothesis of ecological additivity (Huynh *et al*., 2020). Consequently, only deviations beyond ecological additivity should be interpreted as evidence for niche divergence (Parisod & Broennimann, 2016). Such deviations can be decomposed into components of niche stability (S), unfilling (U), and expansion (E) (Guisan et al., 2014), allowing the distinction among niche contraction, dominance, and true niche novelty (Parisod & Broennimann, 2016).

Despite the conceptual importance, explicit tests of ecological additivity that incorporate both parental taxa remain scarce. Robust assessment of niche differentiation requires the inclusion of both parental species (Huynh *et al*., 2020), alongside consideration of environmental conditions across different spatial scales (Leempoel *et al*., 2015; Parisod & Broennimann, 2016). Although coarse-grained climatic variables are widely employed in niche analyses, they are not necessarily representative of the microenvironment experienced by plants, and fine-grained environmental data derived from vegetation records can provide additional resolution (Scherrer & Körner, 2010; Kirchheimer *et al*., 2016; Gorospe *et al*., 2025). Ecological indicator values (EIVs) offer an effective means for characterising local environmental conditions and have been successfully applied in polyploid complexes (Hülber *et al*., 2015; Kirchheimer *et al*., 2016; Silbernagl & Schönswetter, 2019). While direct environmental measurements provide higher resolution (Schaffers & Sýkora, 2000; Wamelink *et al*., 2005), their logistic demands render EIVs a valuable alternative for large-scale studies (Diekmann, 2003; Descombes *et al*., 2020). Ecological divergence may manifest differently across spatial scales. Niches inferred from coarse-grained climatic and edaphic data capture broad environmental tolerances, whereas fine-grained ecological differentiation can reflect local habitat filtering, biotic interactions, and realised niche partitioning, particularly in highly heterogeneous landscapes (Glennon *et al*., 2014; Kirchheimer *et al*., 2016). Failure to distinguish between these grains of analyses risks conflating niche conservatism at a larger scale with ecological divergence at a local scale.

A particularly intriguing case for studying the ecological consequences of hybridisation is *Luzula alpina* (ALP), a tetraploid interploidal hybrid species derived from diploid *L. exspectata* (EXS) and allotetraploid *L. multiflora* (MUL; Heimer *et al*., 2025). Despite the strong reproductive barrier caused by WGD, cross-ploidy hybridisation – particularly involving allopolyploids – may be more common than previously assumed (Abbott & Lowe, 2004; Vallejo-Marin, 2012; Brown *et al*., 2024). Although an increasing number of studies have provided evidence of gene flow between cytotypes (Monnahan *et al*., 2019; Cheng *et al*., 2019; Leal *et al*., 2024), the ecological consequences of such cross-ploidy hybridisation remain poorly understood. As a tetraploid interploidal hybrid, ALP falls conceptually between HHS and allopolyploid speciation, depending on the parental species to which it is compared. Nevertheless, the null hypothesis of genetic – and hence ecological – additivity also applies in this case. Accordingly, only niche shifts beyond parental additivity may be attributed to ecological divergence, that may either be associated with WGD, when compared with EXS, or independent of ploidy, when compared with MUL. To date, only a single study has examined niche differentiation in this group (Geurden *et al*., 2025), reporting limited niche divergence between ALP and EXS. However, this study was restricted to three localities and did not include tetraploid MUL, precluding tests of ecological additivity.

Here, we integrate genomic and environmental data with plot-level vegetation relevés to investigate the ecological consequences of cross-ploidy hybridisation in *Luzula* across the Eastern Alps. Specifically, we (i) reassess the hybrid origin of ALP using double-digest restriction site-associated DNA sequencing (ddRADseq), (ii) characterise population structure, spatial range expansion, and potential glacial refugia for EXS, MUL, and ALP, and (iii) test the null hypothesis of ecological additivity at two spatial resolutions using coarse-grained climatic, topographic, and edaphic data, alongside fine-grained plot-level data based on vegetation relevés and EIVs. Finally (iv), we evaluate morphological differentiation and assess whether ALP exhibits morphological intermediacy relative to its progenitors.

## Materials and Methods

### Study group and population sampling

The genus *Luzula* (Juncaceae) is particularly diverse in the European Alps (Aeschimann *et al*., 2004). Our three study species belong to *Luzula* sect. *Luzula*, which is taxonomically challenging and characterised by extensive hybridisation and allopolyploidisation (Heimer *et al*., 2025; Carrizo García *et al*., 2026), resulting in pronounced morphological homogeneity and frequent misidentifications (Bačič *et al*., 2019). Although EXS and ALP have been reported from the Pyrenees in addition to the Alps (Kirschner, 1996; Carrizo García *et al*., 2026), the precise distributions of our study species have been delimited based on genomic data only for the Eastern Alps (Heimer *et al*., 2025), and we therefore focused on this region. Whereas the ranges of EXS and the hybrid ALP overlap across the Eastern Alps, MUL is parapatric to ALP and limited to the easternmost Alps.

We sampled 53, 20, and 59 populations of EXS, MUL, and ALP, respectively, collecting on average four individuals per population, resulting in 122, 86 and 311 accessions per species (519 in total; Table S1). Leaves for DNA extraction were dried in silica gel and herbarium vouchers were prepared and deposited at IB. Three accessions of *L. campestris* were included as an outgroup. GPS coordinates were recorded for each individual, and mean coordinates were calculated to represent population locations.

### ddRAD sequencing data, alignment and variant calling

Double-digest RADseq data were obtained from Heimer *et al*. (2025). Reads were aligned to the reference genome of closely related *L. pallescens* (NCBI accession GCA_964274315.1) using BWA-MEM v.0.7.17 (Li & Durbin, 2009) with default settings. BAM files were sorted by genomic coordinates, indexed using samtools v.1.9 and read groups were added in Picard v.2.26.2 (http://broadinstitute.github.io/picard/). Single nucleotide polymorphisms (SNPs) were called in GATK v.4.6.2.0 (McKenna *et al*., 2010). *HaplotypeCaller* was run in gVCF for each sample individually, with the correct ploidy specified as inferred from relative genome size estimation (Heimer *et al*., 2025). Following joint genotyping for all samples with *GenotypeGVCF*, biallelic SNPs were selected using *SelectVariants* and filtered according to the Broad Institute’s hard filtering recommendations in *VariantFiltration* (Table S2). We retained only SNPs with a minimum read depth of eight and a maximum read depth of 200.

Two samples with > 50% missing data were excluded from downstream analyses.

### Interspecific genetic differentiation

We first investigated the genomic differentiation among our three study species by constructing a NeighbourNet phylogenetic network in SplitsTree4 (Huson & Bryant, 2006) based on Nei’s distances (Nei, 1972) computed from SNPs present in at least 50% of samples using *StAMPP* (Pembleton *et al*., 2013) in R v.4.5.1 (R Core Team, 2025). The same dataset was also used to perform a principal component analysis (PCA) of genetic variation in *adegenet* (Jombart & Ahmed, 2011). Admixture among the three species was assessed by performing Bayesian clustering in STRUCTURE v.2.3.4 (Pritchard *et al*., 2000), which is robust towards ploidy (Stift *et al*., 2019). The analysis was based on SNPs with a minimum minor allele count (MAC) of three, present in at least 80% of individuals. Only a single SNP per RAD locus was retained by sampling SNPs at least 270 bp from each other to reduce linkage, resulting in 2070 SNPs. STRUCTURE was run with the admixture model for 2,000,000 MCMC generations with 200,000 generations as burn-in for *K* (number of groups) ranging from 1 to 12 with 10 replicates each. STRUCTURE results were aligned across replicates in CLUMPAK (Kopelman *et al*., 2015) and the optimal *K* was determined following Evanno *et al*. (2005) in the R package *pophelper* (Francis, 2017).

We tested the proposed hybrid origin of ALP by computing a hybrid index (Buerkle, 2005) and interspecific heterozygosity based on unadmixed (admixture proportion < 5%) accessions of parental taxa. The hybrid index ranges from zero to one, representing pure individuals of each parental lineage, i.e., EXS and MUL, respectively. A maximum likelihood estimate of the hybrid index and interspecific heterozygosity were computed in *Introgress* (Gompert & Buerkle, 2010) based on a subset of 2096 linkage-pruned SNPs that differed between parental taxa. Because *Introgress* does not accept polyploid genotypes, these were converted into diploid ones by discarding dosage information but preserving homozygote and heterozygote states.

### Population structure and range expansion

Intraspecific population structure was assessed for all three species individually by means of NeighborNets and Bayesian Clustering as described in the previous section. Population differentiation was assessed with Rho, an equivalent of F_ST_ suitable for polyploids (Ronfort *et al*., 1998). Nucleotide diversity (π) was computed in piawka (Scott *et al*., 2025) for all populations with n ≥ 2 sampled individuals from linkage-pruned SNPs present in at least 80% of individuals of the respective species. The same dataset was used to calculate the number of private alleles for each population. Spatial patterns of range expansion were inferred from the directionality index ψ (Peter & Slatkin, 2013, 2015), which leverages differences in the strength and directionality of genetic clines in the allele frequency spectrum to detect the geographic origin and direction of range expansion. Because many Alpine species survived the last glacial period in multiple refugia (Schönswetter *et al*., 2002, 2004), we computed ψ for each major genetic group defined by STRUCTURE in addition to the entire species. We followed Marske *et al*. (2020) to compute ψ with the *X-Origin* pipeline (He *et al*., 2017), using the outgroup *L. campestris* to infer ancestral allelic states. As for the hybrid index, polyploid genotypes were converted into diploid ones for the calculation of ψ. We further investigated whether the inferred origins of range expansion were supported by spatial patterns of nucleotide diversity using linear models to test for the effect of Euclidean distance to the expansion origin on π. The influence of demographic and ecological processes on the spatial structure of genetic variation was assessed by computing isolation-by-distance (IBD) and isolation-by-environment (IBE) within each species (Methods S1).

### Environmental niche modelling

Ecological differentiation among our study species was investigated at two spatial resolutions, using coarse-grained environmental data from publicly available databases and fine-grained data derived from a large set of vegetation plots. We opted for this two-scale approach to rigorously assess differences among study species in abiotic climatic and edaphic niches on the one hand, and microhabitat preferences that are shaped by biotic interactions, on the other hand. Niche differentiation based on the coarse-grained dataset was assessed by constructing environmental niche models (ENMs) and performing tests for niche divergence based on climatic, topographic, lithologic and edaphic variables obtained from public databases (Methods S2).

For ENMs, we selected five biologically relevant environmental predictors representing temperature, precipitation and geology (bio1, bio8, bio12, bio15, bedrock acidity; Table S3), that were only weakly correlated (Spearman’s | r | < 0.7) based on a random sample of 10,000 background points within a buffer of 150 km around occurrences (Dormann *et al*., 2013). A total of 53, 20, and 59 occurrence points were retained for EXS, MUL and ALP, respectively, which were complemented with 1000 pseudo-absences sampled randomly from the background area. Because of the high morphological similarity of the three species and the resulting lack of publicly available and reliable species-specific occurrence data, we were limited to the populations genotyped by us. Accounting for this scarcity in presences, we used ensembles of small models (ESMs; Breiner *et al*., 2015) in *ecospat* (Di Cola *et al*., 2017), which combine many bivariate models into a final model prediction and have proven to be particularly powerful with few occurrence records. ESMs were constructed with four algorithms, each run with 10 replicates: generalised linear models, maximum entropy models (Phillips *et al*., 2006), artificial neural networks (Ripley, 1996), random forest (Breiman, 2001) and generalised additive models (Hastie & Tibshirani, 1986). Individual models with an area under the receiver operating characteristic curve (AUC) > 0.5 were retained and weighted by their AUC to construct the final ESM. Model predictions were converted into binary presence–absence classes using a threshold that maximised the true skill statistic (TSS; Allouche *et al*., 2006). To identify potential glacial refugia, models were projected to climatic conditions during the Last Glacial Maximum (LGM; ca. 21 kya), based on climate data retrieved from CHELSA. Bedrock acidity was considered constant through time. We used four general circulation models from different families to account for uncertainties in paleoclimatic reconstruction (Methods S2; Varela *et al*., 2015) and retained predicted presences only if supported by at least two models.

### Coarse-grained niche divergence

We tested for niche divergence among study species using all available 32 climatic, lithologic, topographic and edaphic variables of the coarse-grained dataset (Table S3). Kernel-smoothed occurrence densities were computed along principal components from a common environmental space gridded with 100 × 100 cells using *ecospat*. Niche overlap was characterised with Schoener’s *D* (Schoener, 1968), ranging from 0 (fully divergent niches) to 1 (completely overlapping niches). We used niche equivalency tests with 1000 replicates each to test if two niches are significantly different (i.e., rejection of niche equivalency if *P* < 0.05). We further evaluated niche overlap with similarity tests, which assess whether the niche of one species predicts the other better than expected by chance. If the observed *D* is lower (*P* < 0.05) than the null distribution, niches are less similar than expected by chance, whereas *D* greater than the null distribution (*P* > 0.95) indicates higher than random similarity (Molina-Henao & Hopkins, 2019).

In addition to pairwise comparisons, we compared the niche of ALP to the combined niche of EXS and MUL to test the null hypothesis of niche additivity (Parisod & Broennimann, 2016) in the hybrid species. Deviations from this null hypothesis were decomposed into normalised components of unfilling, stability, and expansion following the USE framework of Guisan *et al*. (2014). Niche stability S corresponds to the fraction of the observed hybrid niche overlapping with the combined niche of both progenitors. The fraction of the hybrid niche deviating from the additivity of progenitors (i.e. niche shift) was divided into niche contraction U (i.e. the proportion of unoccupied niche space) and niche expansion E (i.e. the proportion of newly occupied niche space).

Niche breadth of each species was computed as the standard deviation of PCA scores along the first two environmental axes and niche optima were inferred from the mean of these scores. Following Theodoridis *et al*. (2013) and Kirchheimer *et al*. (2016), we randomly sampled 100 cells of the gridded environmental space weighted by the density of species occurrences. We repeated this process 1000 times and assessed differences in niche optima and niche breadths using an ANOVA and Tukey HSD *post hoc* test.

The most relevant variables for niche divergence were identified using redundancy analysis (RDA) in *vegan* (Oksanen *et al*., 2025), using occurrences of the three *Luzula* species as response variables and environmental predictors as explanatory variables. An optimised RDA model was built using step-wise model selection based on the Akaike Information Criterion (AIC). The goodness of fit between the optimised and full models was compared with adjusted *R^2^*, and the significance of the entire optimised model, each RDA axis and each explanatory variable was determined by analysis of variance (ANOVA).

### Fine-grained niche divergence

We used vegetation relevés to characterise differences in plant community composition among our study species, and employed EIVs to test niche divergence based on fine-grained environmental data recorded at the plot scale. For each sampled *Luzula* specimen, we registered all vascular plant species and their estimated percentage cover within a radius of 20 cm, as well as additional ecologically relevant variables such as the total plant cover, the cover of dead organic material, rocks, soil and cryptogams (Table S4, S5). Plants were identified using regional floras (Aeschimann *et al*., 2004; Fischer *et al*., 2008; Eggenberg *et al*., 2020), following the nomenclature of Karrer (2024). Elevation was retrieved from a digital elevation model with a resolution of 30 m (Methods S2). Vegetation data were available for 515 genotyped *Luzula* specimens, of which we excluded seven putative backcrosses between ALP and MUL (see Results), two individuals with > 50% missing data in the RADseq dataset and one sample with a strongly divergent plant community, for a final dataset of 505 relevés. Differences in plant community composition were inferred from a Two-way Indicator Species Analysis (TWINSPAN; Hill, 1979) and a detrended correspondence analysis (DCA), which were computed in the R packages *twinspan* (https://github.com/jarioksa/twinspan) and *vegan*, respectively. The significance of interspecific differentiation in the surrounding vegetation composition was determined with an RDA using the presence of the three *Luzula* species as response and the Hellinger-transformed vegetation data as explanatory variables.

Microhabitat preferences of our study species were inferred from community weighted means of Karrer indicator values (Karrer, 2024). We chose Karrer values because they were designed specifically for Austria, which encompasses most of our study region. We discarded three species groups without available Karrer EIVs (2.8% of data) and computed rescaled Landolt EIVs (Landolt, 2010) for four species not occurring in Austria (0.1% of data; Table S6). Species that are indifferent towards a certain EIV were treated as “NA” for that indicator. Analogous to the analysis of the coarse-grained dataset, we tested niche divergence between *Luzula* species in *ecospat* using mean EIVs and additional plot-scale environmental predictors and decomposed niche shifts using the USE framework. Following Geurden *et al*. (2025), we defined the background as the total environmental space occupied by the three species.

Finally, the significance of interspecific differentiation regarding EIVs and plant community composition was assessed by performing an RDA for each of the two data sets, using the three *Luzula* species as response variables and either the fine-grained ecological variables or the surrounding plant species as explanatory variables (Gorospe *et al*., 2025). An optimised RDA model was built and evaluated as described for the coarse-grained dataset and mean differences of ecological variables among study species were assessed by Kruskal-Wallis and Dunn’s *post hoc* test using the R package *rstatix* (Kassambara, 2023).

### Analyses of morphometric data

Morphological differentiation among our study species was assessed through morphometric analyses. A set of 17 characters, including reportedly discriminative ones (Bačič *et al*., 2007; Fischer *et al*., 2008), were measured for 92, 49 and 201 genotyped herbarium specimens of EXS, MUL and ALP, respectively, totalling 342 accessions (Table S7). Seed characters were developed in 292 individuals and analysed separately. Flower and seed characters, as well as the number of teeth on the margin of cauline leaves were measured on images taken with a stereomicroscope Olympus SZX9 using the Olympus image analysis software analySIS pro. In addition to directly measured characters, we calculated six ratios.

Mean differences in morphological traits were assessed through Kruskal-Wallis test followed by Dunn’s *post hoc* test. Missing values were substituted with the mean values of the respective species for multivariate analyses. All characters were weakly correlated (Spearman’s | r | < 0.8) and, after standardisation to zero mean and one unit variance, were used to perform a PCA and linear discriminant analysis (LDA) to explore morphological differentiation and to find the characters best separating the species. PCA and LDA were computed using the packages *MASS* (Venables & Ripley, 2002) and *FactoMineR* (Lê *et al*., 2008), respectively, and were performed separately for seed and all other characters. Accuracy of the LDA was evaluated using leave-one-out cross validation.

## Results

### Interspecific genomic differentiation

EXS, MUL and ALP were clearly separated in the PCA of genomic variation (Fig. 1a) and the NeighborNet (Fig. S1) derived from RADseq data, with the hybrid ALP occupying an intermediate position between the parental species. Accordingly, the average hybrid index of ALP was 0.51 (SD: 0.04, Fig. 1b) and its average interspecific heterozygosity was 0.37 (SD: 0.04). The optimal number of clusters in the STRUCTURE analysis was three (*K* = 3) based on Δ*K* (Figs. 1c, S2, S3), corresponding to the three species. Notably, seven tetraploid individuals were strongly admixed between the genetic clusters of ALP and MUL, occupied intermediate positions between the two species in the PCA and had an elevated hybrid index of 0.74 ± 0.07.

**Fig. 1.**
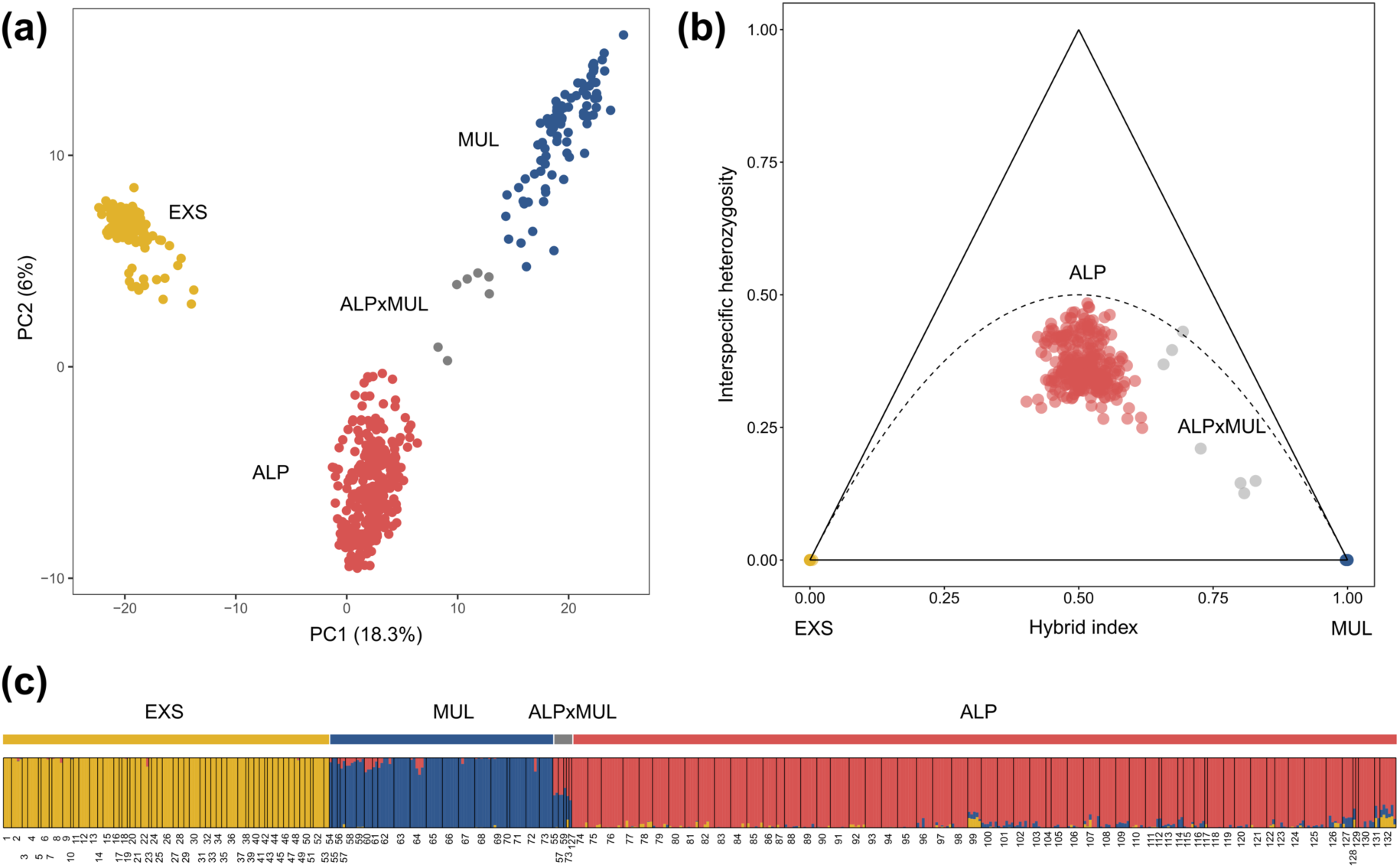
Genetic differentiation among the study species. (a) Principal component analysis (PCA) of genetic variation among *Luzula exspectata* (EXS; yellow), *L. multiflora* (MUL; blue) and *L. alpina* (ALP; red) based on 49,884 SNPs. Seven individuals identified as backcrosses between MUL and ALP are shown in grey. The same colour scheme is used throughout the manuscript. (b) Hybrid index and interspecific heterozygosity calculated from 2096 unlinked SNPs that differed between unadmixed individuals of parental taxa (EXS and MUL). The dotted line represents the expected position of F2 hybrids. (c) Assignment of individuals to genetic clusters inferred by STRUCTURE for the most likely *K* = 3 based on 2070 unlinked SNPs. Numbers indicate populations; species are shown as coloured bars on top, with the seven putative backcrosses between ALP and MUL indicated in grey.

**Fig. 2.**
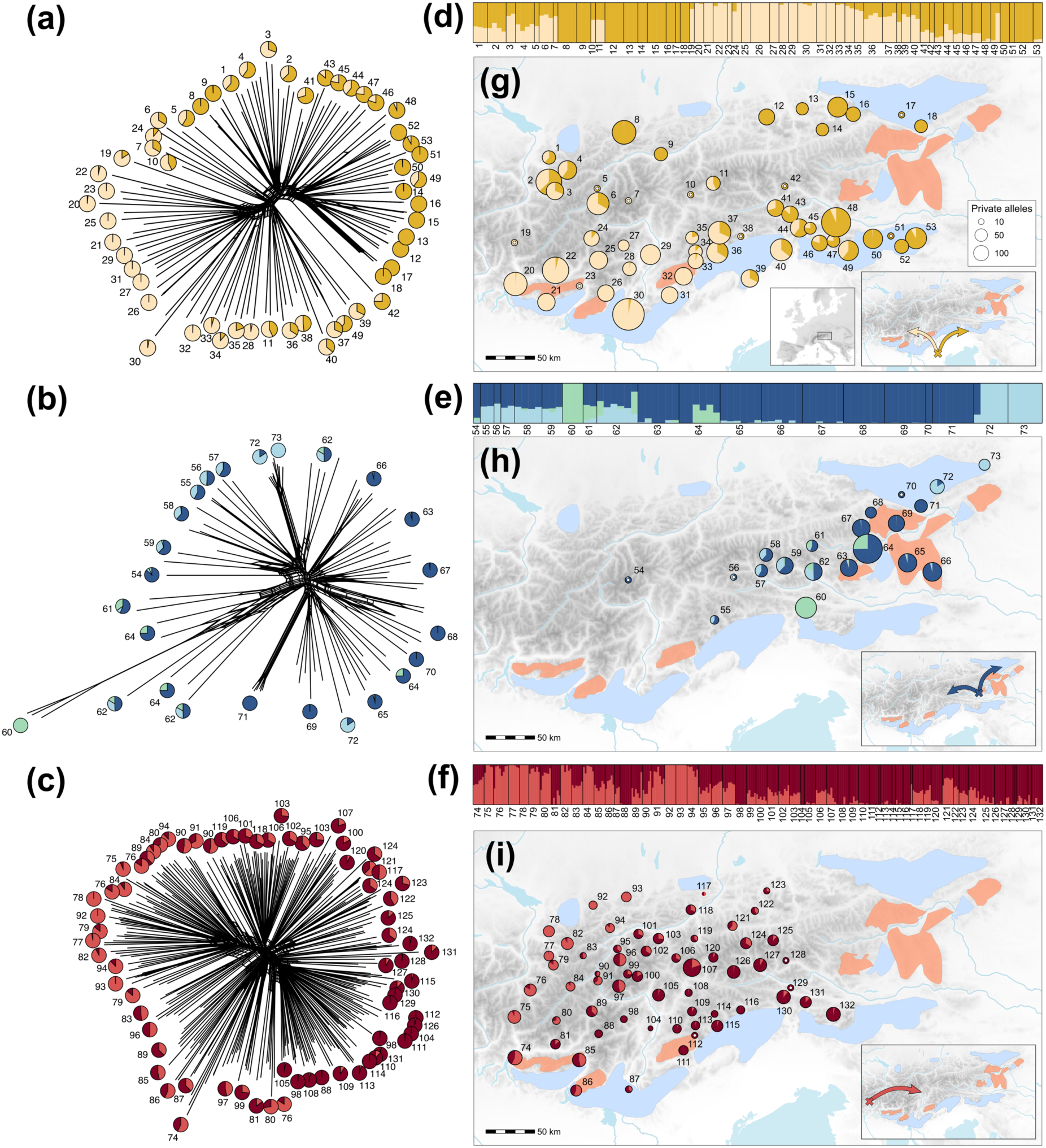
Population structure of *Luzula exspectata* (EXS; top), *L. multiflora* (MUL; middle) and *L. alpina* (ALP; bottom). (a)–(c) NeighbourNets with population-average genetic cluster assignment in STRUCTURE. (d)–(f) Barplots of per-individual genetic cluster assignment in STRUCTURE. (g)–(i) Distribution of population-average genetic cluster assignment. Sizes of pie charts in the maps are proportional to the number of private alleles for populations with n ≥ 2 individuals sampled. Populations with only a single genotyped individual are indicated by white points. Crosses in map inserts in (g)–(i) show the most likely origin of range expansion for the entire species inferred from the directionality index and arrows the hypothesised approximate direction of range expansion. Polygons in light blue and red show calcareous and siliceous potential refugia, respectively, according to Tribsch and Schönswetter (2003).

**Fig. 3.**
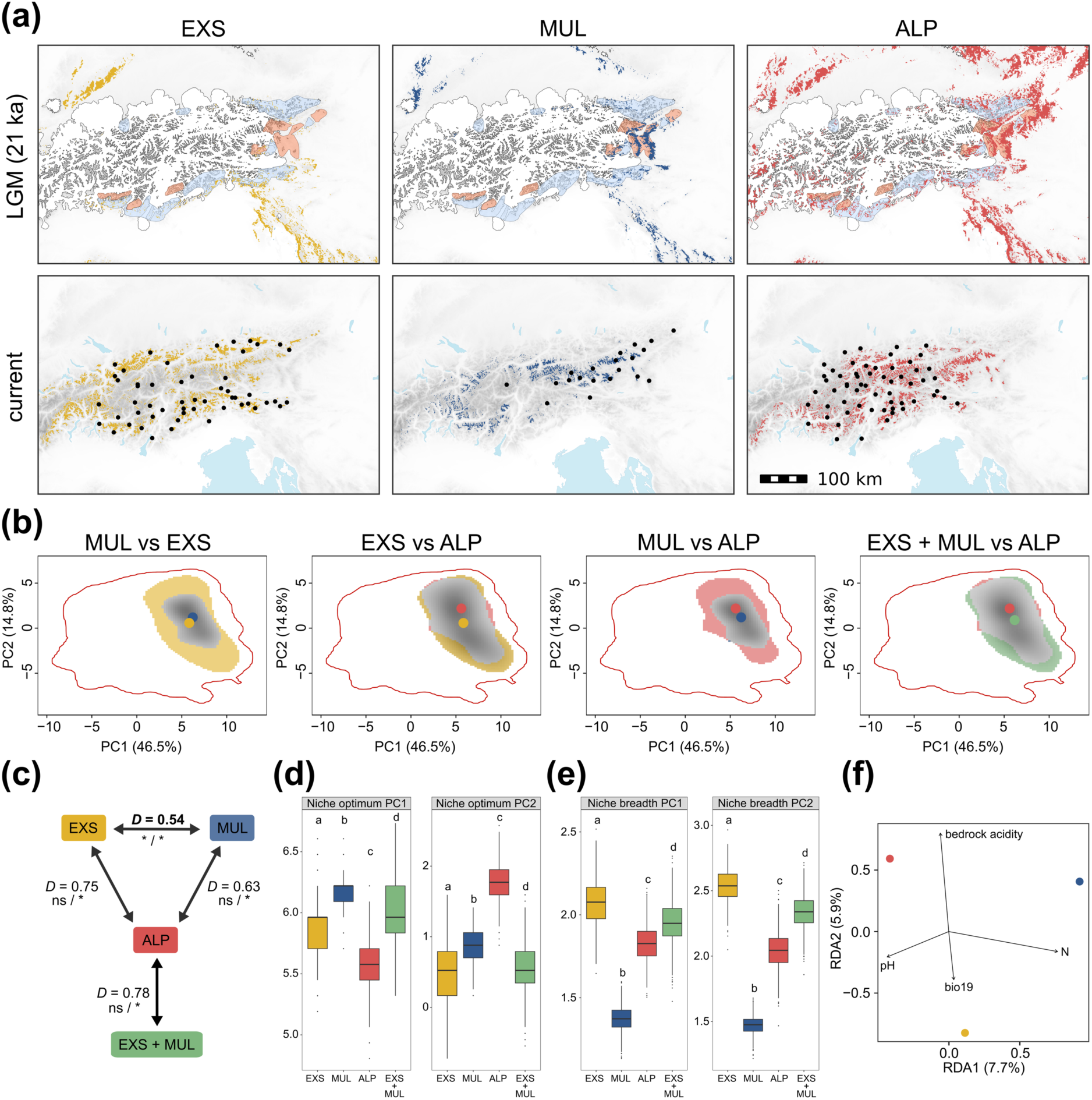
Analysis of coarse-grained environmental data. (a) Predicted suitable area during the Last Glacial Maximum (LGM; top row) and under current climatic conditions (bottom) based on environmental niche modelling for *Luzula exspectata* (EXS), *L. multiflora* (MUL) and *L. alpina* (ALP). Polygons in light blue and red are calcareous and siliceous potential refugia, respectively, according to Tribsch and Schönswetter (2003). The extent of the LGM ice sheet is shown in white following Ehlers *et al*. (2011) and van Husen (1987). Black dots in the “current” panel are occurrences used for model parameterisation. (b) Pairwise comparison of ecological niches. ALP was additionally compared to the combined niche of its progenitors (EXS + MUL) to test for deviations from the expected ecological additivity. Coloured regions show ecological niches for each species along the first two principal components of multivariate niche space while overlapping niches are shown in grey. The background area is indicated by a red line and coloured dots are niche centroids. (c) Schoener’s *D* for each pairwise comparison of niche overlap between species. Bold font indicates significantly non-identical niches. The significance of niche equivalency and similarity tests is indicated below (ns: not significant, * *P* < 0.05). (d) Niche optima and (e) niche breadths along the first two principal components of environmental space. Letters in (d) and (e) indicate significant (*P* < 0.05) differences according to Tukey’s HSD *post hoc* test. (f) Optimised redundancy analysis (RDA) showing the four most relevant coarse-grained environmental variables for ecological differentiation among study species.

### Population structure and range expansion

Geographic patterns of genetic variation were weak for all three species, as reflected by star-like NeighborNets (Fig. 2a–c). Genetic differentiation among populations was on average highest for EXS (Rho = 0.649; Fig. S4), followed by MUL (0.406; Fig. S5) and ALP (0.354; Fig. S6) and was driven by IBD (Tables S8, S9, Fig. S7).

Signatures of IBE were significant across analyses only for the hybrid ALP and were supported by RDA but not by Mantel tests for EXS (Tables S8, S9, Fig. S8).

The optimal number of clusters was *K* = 2 for both EXS and ALP (Fig. S2), roughly corresponding to a northwest-southeast and southwest-northeast cline of genetic variation, respectively (Fig. 2d,f,g,i). At the best *K* = 3, populations of MUL segregated into one genetic cluster predominantly in the easternmost Central Alps, another cluster to the west and northeast of this region and one divergent population in the Southern Alps (Fig. 2e,h). Admixture among genetic clusters was present for most populations of MUL. For the other two species, which had a stronger geographic segregation of genetic clusters, admixture was observed in populations where these clusters met (Figs. 2, S9–S11).

The average number of private alleles was highest in EXS (Fig. 2g). Generally, the number of private alleles (nPA) was highest in the Southern Alps, particularly in populations 30 and 48. MUL had the second highest nPA on average, with population 64 ranking first (Fig. 2h). The same population also had the highest π (0.177). There was no apparent pattern in the distribution of the overall fewer private alleles in ALP (Fig. 2i). Tests for range expansion using allele frequency patterns (ψ) revealed significant (*P* < 0.05) expansion with species-specific spatial dynamics. The most likely origin of range expansion for EXS was to the south of the Alps (Fig. 2g), whereas for MUL, the origin coincided with the western-most glacial refugium on siliceous bedrock in the eastern Central Alps, which is still inhabited by the species (Fig. 2h). Lastly, an eastward expansion, possibly from the Western Alps, was inferred for ALP (Fig. 2i). Testing the origin of largely parapatric genetic clusters at *K* = 2 for EXS and ALP suggested similar origins as for the entire species (Fig. S12). Nucleotide diversity π decreased significantly (*P* < 0.05) with geographic distance from the inferred origin of range expansion for all species (Fig. S13; Tables S10, S11).

### Environmental niche modelling

All modelling algorithms and ensemble models showed a good performance (AUC > 0.8). Niche projections were largely congruent with the current distributions of our study species in the Eastern Alps (Fig. 3a). However, some suitable areas in the easternmost Alps were inferred for ALP, whereas our extensive sampling suggests that the species is absent in this area. Conversely, small areas of suitable habitat towards the west were suggested for MUL in a region to our knowledge occupied only by ALP. Projections of occurrence probability to LGM conditions suggest different refugia for the three species. In addition to the northern Dinaric Mountains to the southeast and the Swabian Jura to the northwest of the Eastern Alps, suitable habitat for EXS was inferred in the Southern Alps, a region that acted as important glacial refugium for calcicolous species (Fig. 3a). Model predictions suggest a survival of MUL during the LGM in unglaciated regions of the easternmost siliceous Central Alps. The same region was inferred as potential refugium for ALP, although several regions in the Northern and Southern Alps as well as at the periphery of the Alps (Swabian Jura, Dinaric Mountains, Bohemian Massif) may have also provided suitable conditions.

### Coarse-grained niche divergence

The environmental niches of EXS and MUL showed limited overlap (Schoener’s *D* = 0.54) and were not equivalent (*P* < 0.05; Figs. 3b,c, S14; Table S12). The niche of the hybrid species ALP strongly overlapped with that of MUL (*D* = 0.63) and EXS (*D* = 0.75) and niche equivalency could not be rejected (*P* > 0.05). Similarly, niche overlap was strong between ALP and the combined niche of its parental species (*D* = 0.78, *P* > 0.05), suggesting ecological additivity. Niche conservatism was supported by similarity tests (*P* < 0.05) for all comparisons. Accordingly, when decomposing niche overlap following the USE framework, we found a high degree of niche stability (S = 0.99) between the hybrid species ALP and the combined niche of its progenitors, with only limited evidence of niche unfilling (U = 0.10) and no signal of expansion (E = 0.00). Despite strongly conserved niches, niche optima differed significantly among species along the first two principal components (Fig. 3d).

Corresponding to ecological intermediacy of the hybrid, the niche breadth of ALP was between that of EXS and MUL along both PCs (Fig. 4e) but significantly narrower than the combined niche of both parental species, consistent with niche unfilling.

**Fig 4.**
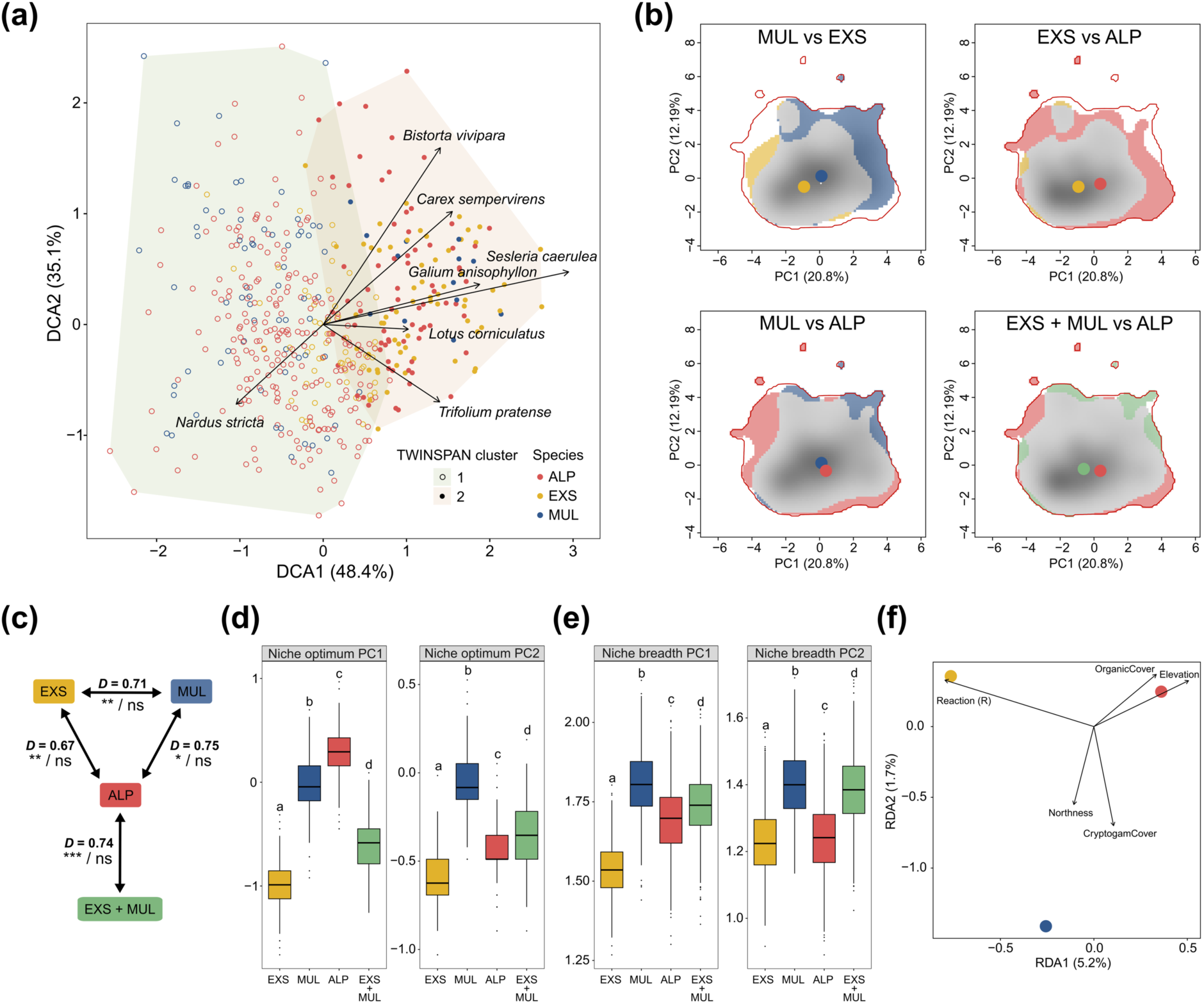
Analysis of fine-grained environmental data. (a) Detrended correspondence analysis (DCA) of plant community composition across relevés coloured by *Luzula* species. Coloured convex-hulls and filled or unfilled circles indicate groups according to the first-level TWINSPAN clustering. Arrows represent accompanying plant species that differentiate these clusters and are typical for species-poor acidic alpine meadows and alpine grasslands on calcareous substrates for clusters one and two, respectively. (b) Pairwise comparison of ecological niches derived from fine-grained ecological data. ALP was additionally compared to the combined niche of its progenitors (EXS + MUL) to test for deviations from the expected ecological additivity. Coloured regions show ecological niches for each species along the first two principal components of multivariate niche space while overlapping niches are shown in grey. The background area is indicated by a red line and coloured dots are niche centroids. (c) Schoener’s *D* for each pairwise comparison of niche overlap between species. Bold font indicates significantly non-identical niches. The significance of niche equivalency and similarity tests is indicated below (ns: not significant, * *P* < 0.05, ** *P* < 0.01, *** *P* < 0.001). (d) Niche optima and (e) niche breadths along the first two principal components of environmental space. Letters in (d) and (e) indicate significant (*P* < 0.05) differences according to Tukey’s HSD *post hoc* test. (f) Optimised RDA of fine-grained ecological variables most differentiating between the study species.

Variables underlying divergent niche optima were identified by an optimised RDA model (adjusted *R^2^* = 0.19), which significantly (*P* ≤ 0.01) explained variation in the data (Table S13). Differences in individual niches were mostly driven by bedrock acidity, topsoil pH and nitrogen content, as well as by the precipitation of the coldest quarter (bio19; Fig. 3f).

### Fine-grained niche divergence

We recorded a total of 393 accompanying vascular plant species (Table S4). With a median of n = 14 plant species per relevé, species richness was highest for EXS, followed by ALP (n = 11) and MUL (n = 10). The communities of accompanying plant species were strongly overlapping among our study species, as reflected by the lack of separation in the DCA and TWINSPAN clusters that did not align with the three *Luzula* species (Fig. 4a; Results S1).

Despite the strong overlap in surrounding plant communities, RDA identified several diagnostic species, which in case of EXS were mostly characteristic for alpine grasslands over limestone, whereas ALP was characterised by the presence of species typical for acidic alpine meadows (Fig. S15). Finally, MUL co-occurred with acidophilic species of alpine dwarf-shrub heaths. The optimised RDA model performed similarly (adjusted *R^2^* = 0.35) to the full model (*R^2^* = 0.38) and the model, both axes and all explanatory variables included were significant (*P* < 0.05; Table S14).

Ecological indicator values were available for 390 (99%) species (Table S6). Fine-grained ecological niches reconstructed from EIVs overlapped strongly between the tetraploids (*D* = 0.75), whereas diploid EXS was slightly more divergent from MUL (*D* = 0.71) and ALP (*D* = 0.67; Figs. 4b,c, S16; Table S15). Niche equivalency was rejected for all pairwise comparisons between species (*P* < 0.05). Although the combined niche of both parental taxa strongly overlapped with the hybrid niche (*D* = 0.74), they were not identical (*P* < 0.05). Niche similarity tests were not significant for any of the comparisons, meaning that one niche explained the other as well as expected by chance, thus supporting neither niche conservatism nor divergence (Table S15).

Decomposing niche overlap between ALP and the combined niche of its progenitors into components of niche unfilling, stability and expansion revealed predominant niche stability (S = 0.98). Only small components of niche unfilling (U = 0.02) and expansion (E = 0.02) were observed, suggesting limited niche evolution. Despite significantly different niche optima (Fig. 4d), the niche breadth of ALP along PC1 and PC2 was intermediate between those of its parental species (Fig. 4e).

The optimised RDA of fine-grained ecological variables significantly explained variation in the data (adjusted *R^2^* = 0.12, *P* < 0.05; Table S16) and identified the EIV for soil reaction (R), elevation, cryptogam cover, dead organic material cover, and northness as the main drivers of differentiation among the study species (Figs. 4f, S17).

### Morphological differentiation

The PCA of non-seed characters revealed strong morphological overlap among the three species (Fig. S18). Nevertheless, a slight separation was observed in the LDA (Fig. 5a), which had a moderate accuracy (71%) and discriminatory power (Wilk’s Lambda = 0.46, *P* < 0.001). ALP occupied an intermediate position between EXS and MUL along LD1. Characters that contributed most to this separation include flower traits (anther length, anther-filament ratio and the width of the perigon leaf margin) as well as the width of basal leaves and number of teeth per mm leaf margin of cauline leaves (Fig. 5b,c, S19). Despite significant differences in the length of seeds and elaiosomes (Fig. 5d, S20), multivariate analyses of seed characters showed strong overlap among species (Figs. S21, S22) and the LDA of seed characters performed poorly (accuracy 55%, Wilk’s Lambda = 0.88, *P* < 0.001).

**Fig 5.**
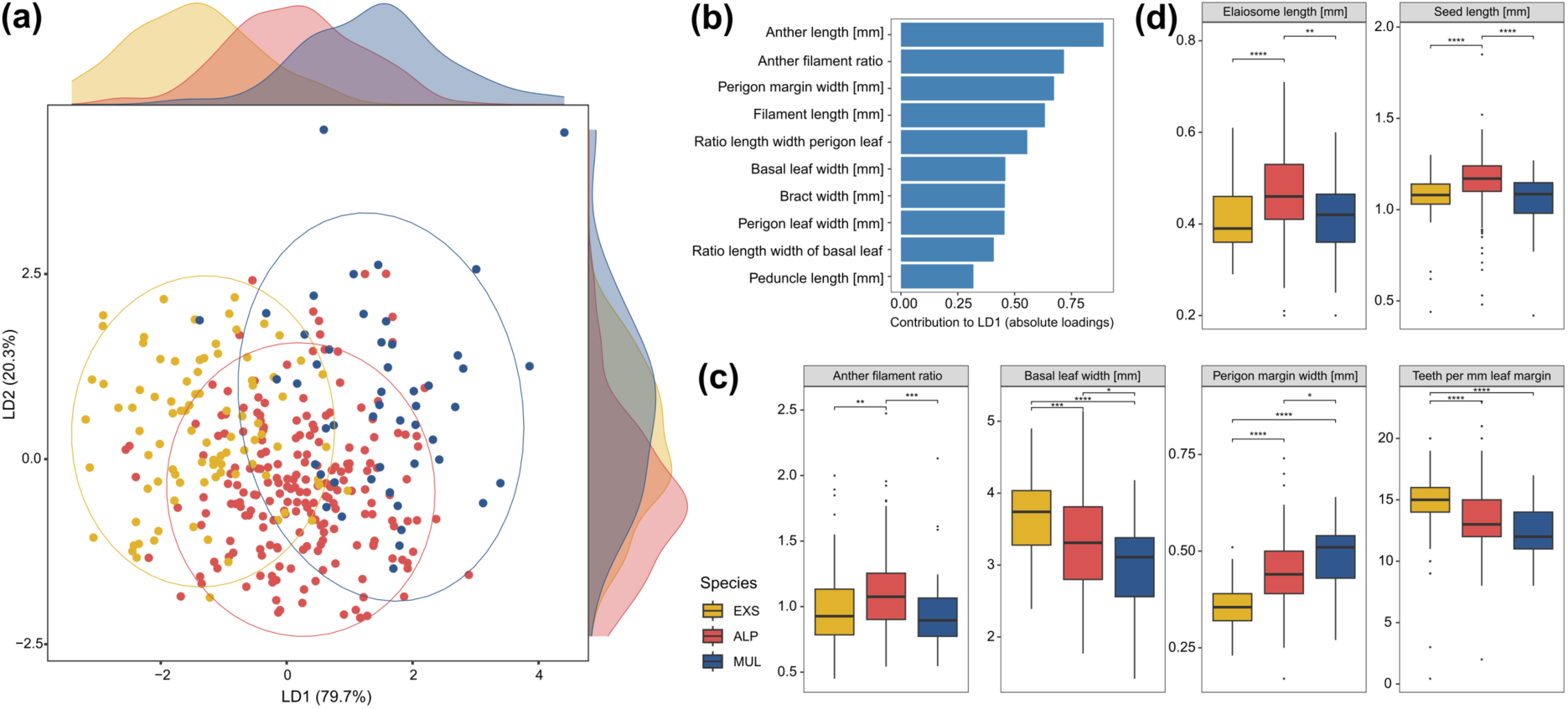
Morphological differentiation among *Luzula exspectata* (EXS; yellow), *L. alpina* (ALP; red) and *L. multiflora* (MUL; blue). (a) Linear discriminant analysis based on 15 morphological characters. Ellipses show 95%-quantiles of samples for each species and marginal densities of occurrences along LD1 and LD2 are shown above and to the right of the plot. (b) Contribution of morphological characters to the first discriminant function in (a) as absolute values of variable loadings. (c) Selected non-seed characters and (d) seed characters with significant differences among species. Asterisks in (c) and (d) indicate significance according to Dunn’s *post hoc* test (* *P* < 0.05, ** *P* < 0.01, *** *P* < 0.001).

## Discussion

### *Luzula alpina* is a cross-ploidy hybrid lineage

Despite the strong reproductive isolation typically exerted by differences in ploidy (Levin, 2002; Ramsey & Schemske, 2002), our genomic analyses support the cross-ploidy hybrid origin of tetraploid ALP proposed by Heimer *et al*. (2025). The karyotype of ALP further supports this scenario, consisting of 24 fragmented, half-size chromosomes matching those of EXS, and 12 full-size chromosomes likely originating from MUL (Nordenskiöld, 1951; Kirschner *et al*., 1988; Heimer *et al*., 2025). These findings add to growing evidence that cross-ploidy hybridisation, particularly involving allopolyploids, is more common than previously assumed (Brown *et al*., 2024). Interspecific heterozygosity of ALP was on average lower than expected for F2 hybrids (Fig. 1b), indicating an older origin (De La Torre *et al*., 2015; Wiens *et al*., 2025), a conclusion also supported by the distinct genetic clusters formed by the three species (Figs. 1c, S3).

Postzygotic isolation resulting from ploidy differences plays no role between ALP and its parent MUL, as both are tetraploid. Although hybrid sterility also occurs without ploidy differences (Koide *et al*., 2018) and could be caused by the divergent karyotypes of ALP and MUL, signatures of admixture and putative backcrosses indicate ongoing gene flow and at least partial fertility between ALP and MUL (Figs. 1, S1). This incomplete reproductive isolation is in line with the reported successful meiotic pairing of fragmented and unfragmented chromosomes in *Luzula* (Nordenskiöld, 1956) and suggests that ALP may represent a case of incipient hybrid speciation (Harrison & Harrison, 1993; Suarez-Gonzalez *et al*., 2018; Rosser *et al*., 2024). In contrast, the ploidy difference between ALP and EXS likely provides effective postzygotic isolation, preventing extensive hybridisation and enabling the partially sympatric distributions of both species, as supported by the absence of triploids in localities where both species co-occur (Geurden *et al*., 2025; this study).

The two parental species are almost fully parapatric, with EXS occurring mostly on calcareous bedrock and MUL being restricted to the easternmost siliceous Central Alps. Consistent with the expected genomic and ecological additivity of hybrids, ALP occupies a larger area across the Alps, albeit with a preference for more acidic substrates (Fig. 2, Results S1). Consequently, the three species persisted in distinct refugia corresponding to their ecological preferences during the LGM, from which they re-colonised their current ranges (Figs. 2, 3; Results S2). Coherent with its smaller range and limited post-glacial expansion, MUL was the only species without significant signals of isolation-by-distance. On the other hand, patterns of isolation-by-environment were consistently recovered only for ALP. Although contrasting the expectation of polyploids expanding rapidly across larger areas (Huynh *et al*., 2020; Moura *et al*., 2021), these results may reflect local adaptation to divergent environmental conditions across the larger range of ALP (Funk *et al*., 2011).

### Contrasting patterns of niche breadths across grains of analyses

Niche breadth estimates differed strikingly between the coarse- and fine-grained analyses. Consistent with its restricted distribution, MUL exhibited the narrowest niche of all species inferred from coarse-grained environmental data. In contrast, it showed the broadest niche at the plot scale, indicating effective exploitation of locally available microhabitats, which are particularly abundant in topographically complex mountain areas (Scherrer & Körner, 2010). Conversely, EXS displayed broad climatic tolerances but comparatively narrow plot-level niches. This discrepancy likely reflects the greater environmental heterogeneity captured across the larger geographic range of EXS, which may inflate estimates of niche breadth at coarse resolutions, whereas in reality the species occupies microsites in which environmental conditions are more constant. Similar scale-dependent patterns have been reported previously (Kirchheimer *et al*., 2016) and underscore a general limitation of coarse-grained environmental data. By ignoring microsite conditions and biotic interactions, such data may lead to biased results, particularly in highly heterogeneous ecosystems (Descombes *et al*., 2020).

The niche breadth of the hybrid ALP was intermediate between its parental species across analyses, indicating that the fixed heterozygosity conferred by cross-ploidy hybridisation is not necessarily associated with niche expansion. Although its broader niche compared to EXS at the plot-scale suggests a better exploitation of available microhabitats, ALP does not exhibit an elevated tolerance towards a wider spectrum of environmental conditions compared to its diploid progenitor. These results align with increasing evidence for idiosyncratic niche dynamics following WGD in both auto- (Glennon *et al*., 2014; Grünig *et al*., 2024; Meirmans & Kolář, 2025) and allopolyploids (Blaine Marchant *et al*., 2016; Huynh *et al*., 2020; Akiyama *et al*., 2021). The ecological consequences of WGD thus appear contingent not only on the type of polyploidy but also on additional factors such as the available habitat and time to expand niche breadths (Kirchheimer *et al*., 2016). Moreover, post-WGD evolution and interploidy introgression play an important role in the ecological divergence of polyploids and may confound the inferred effects of WGS when not being accounted for (Padilla-García *et al*., 2023).

### Cross-ploidy hybridisation within a largely conserved niche

Whereas homoploid hybrid lineages often exhibit pronounced niche shifts due to the absence of intrinsic reproductive barriers (Gross & Rieseberg, 2005; Mao & Wang, 2011) allopolyploids frequently display niche conservatism, facilitated by immediate reproductive isolation via WGD (Blaine Marchant *et al*., 2016; Huynh *et al*., 2020). Cross-ploidy hybrids represent an intermediate scenario, for which ecological outcomes remain poorly understood.

Our analyses identify niche stability as the predominant pattern of niche evolution in the cross-ploidy hybrid ALP. Notably, the extent of deviation from the expected ecological additivity depends on the spatial resolution of environmental data. Coarse-grained environmental data supported the null hypothesis of ecological additivity between the hybrid ALP and its progenitors (Fig. 3), similar to the well-studied example *Aegilops* (Huynh *et al*., 2020). In contrast, fine-grained analyses revealed subtle but significant niche differentiation of ALP (Fig. 4). This discrepancy between grains of analyses likely reflects differences in microclimatic conditions and biotic interactions that alter the species’ realised niche at the plot-level and go undetected by coarse climatic and edaphic data (Kirchheimer *et al*., 2016).

However, the magnitude of niche divergence was small even at the plot-level and niche stability was the dominant pattern of niche evolution across analyses. Together with the intermediate niche breadth of the hybrid species and high levels of niche overlap with both progenitors, these results support niche intermediacy (Blaine Marchant *et al*., 2016) and reject the alternative scenarios of niche dominance (i.e., the hybrid niche is fully nested within one progenitor, but does not fill the niche of the other) or niche contraction (i.e., the hybrid partially fills the niche of both progenitors, but has a narrower niche; Parisod & Broennimann, 2016).

Strong niche conservatism between MUL and ALP appears difficult to reconcile with the incomplete reproductive isolation between the hybrid and its progenitor. However, coexistence of cytotypes within overlapping niches has been documented in other systems, including *Arabidopsis arenosa* (Padilla-García *et al*., 2023), in which gene flow occurs between diploids and autotetraploids (Yant *et al*., 2013). Some species escape competition with their progenitors through changes in phenology, pollinators or post-pollination prezygotic barriers (Segraves & Thompson, 1999; Castro *et al*., 2020), but these mechanisms are probably not relevant for wind-pollinated *Luzula* species. Instead, the largely parapatric distribution of ALP and MUL likely reduces direct competition and facilitates persistence despite niche similarity.

### Morphological intermediacy following cross-ploidy hybridisation

Consistent with the reported morphological homogeneity within *Luzula* sect. *Luzula* (Kirschner, 1996; Bačič *et al*., 2019), the morphospaces occupied by our study species overlapped substantially. Nevertheless, multivariate analyses revealed some level of separation among the three species, which was driven by both vegetative and reproductive traits (Fig. 5), partly confirming characters previously suggested to be discriminative between the species (Bačič *et al*., 2007; Fischer *et al*., 2008). Vegetative traits of ALP were largely intermediate between those of EXS and MUL, supporting morphological intermediacy rather than mosaic phenotypes (Rieseberg *et al*., 1993), a pattern also observed in other *Luzula* species and across different plant genera (Carney *et al*., 2000; Zika *et al*., 2015; Pliszko & Kostrakiewicz-Gierałt, 2018).

In contrast, several reproductive traits were transgressively expressed in the hybrid ALP. Although the fitness consequences of these shifts remain uncertain, strong effects appear unlikely in this system, given the close relatedness of the parental species and the positive association between heterosis and parental genetic divergence (East, 1936; Heimer *et al*., 2025). Consistent with this interpretation, niche stability and stronger signals of niche unfilling than expansion suggest no enhanced capacity of ALP to colonise ecologically novel habitats. Rather than being the result of increased ecological vigour, the broad geographic distribution of ALP may instead be linked to fixed heterozygosity in the hybrid polyploid, which masks deleterious recessive alleles, thereby reducing genetic load and inbreeding depression (Soltis & Soltis, 2000). Like other allopolyploids, the species may therefore be more tolerant of recurrent founder events associated with range expansion (Rosche *et al*., 2017).

Together, our results reconcile apparently contrasting signals of niche conservatism and divergence. Cross-ploidy hybridisation in ALP is predominantly associated with niche stability, while subtle ecological differentiation emerges only at fine spatial scales, yet is reflected in species-specific range dynamics. These findings emphasise that integrating environmental data with different spatial resolutions can alter conclusions about hybrid niche evolution and provide empirical evidence that ecological additivity also applies to cross-ploidy hybrids.

## Acknowledgements

This study was supported by the European Region Tyrol-South Tyrol-Trentino (EGTC) through the Euregio Science Fund, project LUZALP [grant number IPN 133, 4th call 2020]. The field work was partly supported by a bilateral cooperation funded by the Slovenian Research Agency (ARRS) and the Austrian Agency for Education and Internationalisation [ÖAD; SI 09/2018]. The authors thank T. Bačič, D. Baumgartner, J. Czogalla, D. Dietrich, F. Faltner, J. Geurden, S. Kleiner, Ž. Lobnik Cimerman, J. Maindok, A. Mayerova, I. Pungaršek, Š. Pungaršek and G. Span for help during the fieldwork. We also thank M. Heck, M. Helling, N. Kremmel, M. Magauer, D. Pirkebner, N. Rainer, U. Schmidt and F. Tilg for their support with laboratory work and T. Zeni for methodological advice and helpful discussions. Lastly, we are grateful to the Research Area Scientific Computing of the University of Innsbruck for providing access to the Leo3e and Leo4 HPC clusters for bioinformatic analyses.

## Author contributions

VH and BF conceived and designed the study and conducted field work. VH conducted laboratory work, analysed genomic and ecological data and wrote the initial version of the manuscript. BF and PS supervised the analyses and the manuscript drafting. All authors read and edited the final version of the manuscript.

## Data availability

Raw data are publicly available at the Sequence Read Archive under the BioProjects PRJNA1313421 and PRJNA1225458. The data and code that support the findings of this study are openly available at the Open Science Framework (OSF) repository at DOI: https://doi.org/10.17605/OSF.IO/6AM4X

## Supporting Information (brief legends)

**Fig. S1** NeighbourNet of the three study species derived from RADseq data

**Fig. S2** Delta K plots of the STRUCTURE analyses

**Fig. S3** Results of the STRUCTURE analysis including all three study species

**Fig. S4** Heatmap of Rho for *Luzula exspectata*

**Fig. S5** Heatmap of Rho for *Luzula multiflora*

**Fig. S6** Heatmap of Rho for *Luzula alpina*

**Fig. S7** Isolation-distance-distance

**Fig. S8** Isolation-distance-environment

**Fig. S9** Results of the STRUCTURE analysis for *Luzula exspectata*

**Fig. S10** Results of the STRUCTURE analysis for *Luzula multiflora*

**Fig. S11** Results of the STRUCTURE analysis for *Luzula alpina*

**Fig. S12** Inferred origins of range expansion

**Fig. S13** Relationship between nucleotide diversity and distance to the expansion origin

**Fig. S14** Results of coarse-grained niche analysis

**Fig. S15** Results of the RDA of vegetation data

**Fig. S16** Results of fine-grained niche analysis

**Fig. S17** Box plots of fine-grained environmental data

**Fig. S18** PCA of non-seed morphological characters

**Fig. S19** Box plots of non-seed morphological characters

**Fig. S20** Box plots of seed morphological characters

**Fig. S21** PCA of seed morphological characters

**Fig. S22** LDA of seed morphological characters

**Table S1** *<u>Luzula</u>* accessions used in this study. This table is available as a separate spreadsheet

**Table S2** Details of filtering applied to ddRADseq data

**Table S3** Coarse-grained data used for environmental niche modelling and niche analysis

**Table S4** Vegetation relevé data. This table is available as a separate spreadsheet

**Table S5** Fine-grained ecological data. This table is available as a separate spreadsheet.

**Table S6** Taxa with no Karrer ecological indicator values available.

**Table S7** Morphometric data. This table is available as a separate spreadsheet.

**Table S8** Results of Mantel tests for isolation-by-distance and isolation-by-environment

**Table S9** Results of RDA for isolation-by-distance and isolation-by-environment

**Table S10** Origins of range expansion

**Table S11** Linear models of nucleotide diversity and distance to the origin of range expansion

**Table S12** Results of coarse-grained niche analysis

**Table S13** RDA of coarse-grained environmental variables

**Table S14** RDA of vegetation data

**Table S15** Results of fine-grained niche analysis

**Table S16** RDA of fine-grained environmental variables

**Methods S1** Isolation-by-distance and isolation-by-environment

**Methods S2** Environmental data acquisition

**Results S1** Analyses of vegetation data

**Results S2** Idiosyncratic patterns of refugial areas and range expansion

