## Supporting Information for "Cross-ploidy hybridisation in Alpine woodrushes is associated with ecological additivity and scale-dependent niche divergence"

New Phytologist

**_New Phytologist_ Supporting Information**

The following Supporting Information is available for this article:

**Fig. S1** NeighbourNet of the three study species derived from RADseq data
**Fig. S2** Delta *K* plots of the STRUCTURE analyses
**Fig. S3** Results of the STRUCTURE analysis including all three study species
**Fig. S4** Heatmap of Rho for *Luzula exspectata*
**Fig. S5** Heatmap of Rho for *Luzula multiflora*
**Fig. S6** Heatmap of Rho for *Luzula alpina*
**Fig. S7** Isolation-distance-distance
**Fig. S8** Isolation-distance-environment
**Fig. S9** Results of the STRUCTURE analysis for *Luzula exspectata*
**Fig. S10** Results of the STRUCTURE analysis for *Luzula multiflora*
**Fig. S11** Results of the STRUCTURE analysis for *Luzula alpina*
**Fig. S12** Inferred origins of range expansion
**Fig. S13** Relationship between nucleotide diversity and distance to the expansion origin
**Fig. S14** Results of coarse-grained niche analysis
**Fig. S15** Results of the RDA of vegetation data
**Fig. S16** Results of fine-grained niche analysis
**Fig. S17** Box plots of fine-grained environmental data
**Fig. S18** PCA of non-seed morphological characters
**Fig. S19** Box plots of non-seed morphological characters
**Fig. S20** Box plots of seed morphological characters
**Fig. S21** PCA of seed morphological characters
**Fig. S22** LDA of seed morphological characters

**Table S1** *Luzula* accessions used in this study. This table is available as a separate spreadsheet
**Table S2** Details of filtering applied to ddRADseq data
**Table S3** Coarse-grained data used for environmental niche modelling and niche analysis
**Table S4** Vegetation relevé data. This table is available as a separate spreadsheet
**Table S5** Fine-grained ecological data. This table is available as a separate spreadsheet.
**Table S6** Taxa with no Karrer ecological indicator values available.
**Table S7** Morphometric data. This table is available as a separate spreadsheet.
**Table S8** Results of Mantel tests for isolation-by-distance and isolation-by-environment
**Table S9** Results of RDA for isolation-by-distance and isolation-by-environment
**Table S10** Origins of range expansion
**Table S11** Linear models of nucleotide diversity and distance to the origin of range expansion
**Table S12** Results of coarse-grained niche analysis

**Table S13** RDA of coarse-grained environmental variables
**Table S14** RDA of vegetation data
**Table S15** Results of fine-grained niche analysis
**Table S16** RDA of fine-grained environmental variables

**Methods S1** Isolation-by-distance and isolation-by-environment
**Methods S2** Environmental data acquisition
**Results S1** Analyses of vegetation data
**Results S2** Idiosyncratic patterns of refugial areas and range expansion

**Fig. S1** NeighbourNet of the three study species based on Nei's distances computed from 49,884 SNPs present in at least 50% of samples. The three species and putative backcrosses (ALPxMUL) between *Luzula alpina* (ALP) and *L. multiflora* (MUL) are indicated with colours.

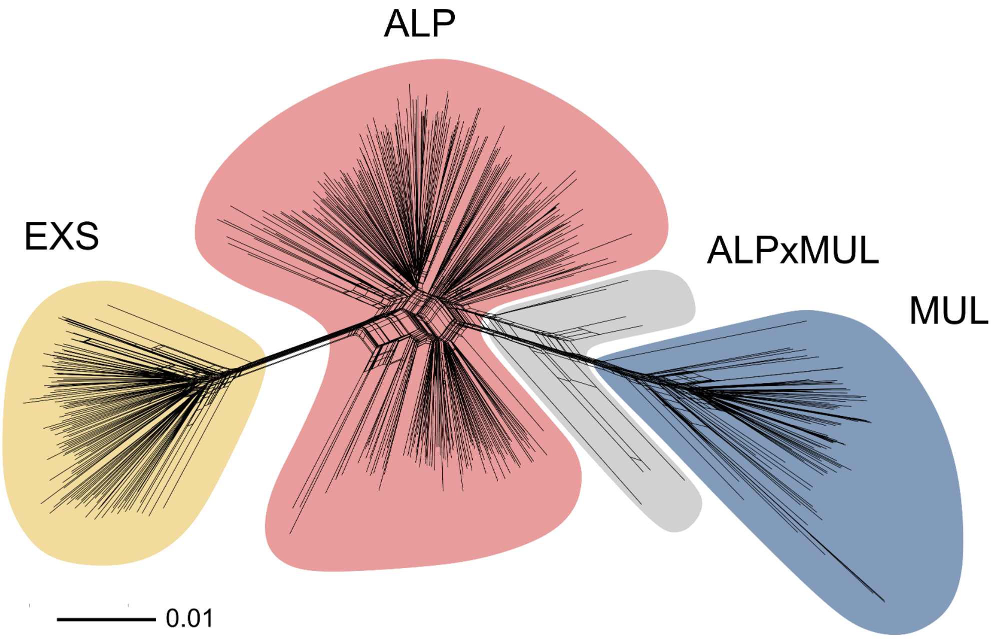


**Fig. S2** Delta *K* plot based on the rate of change in the log probability between successive *K* values in the STRUCTURE analysis for the full data set (a), *Luzula expectata* (EXS; b), *L. multiflora* (MUL; c) and *L. alpina* (ALP; d).

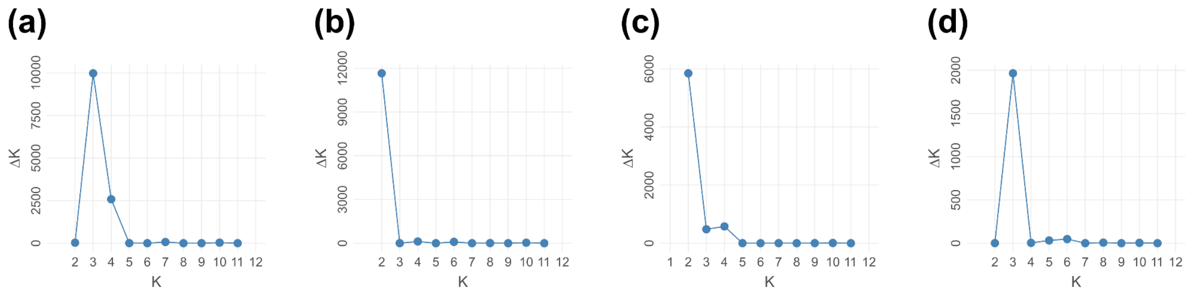

New Phytologist

**Fig. S3** STRUCTURE results for *Luzula exspectata* (EXS), *L. multiflora* (MUL) and *L. alpina* (ALP) for $K$ = 1 to 12. Species and the seven putative backcrosses between ALP and MUL are indicated with coloured bars at the top. Numbers below bar charts are population identifiers.

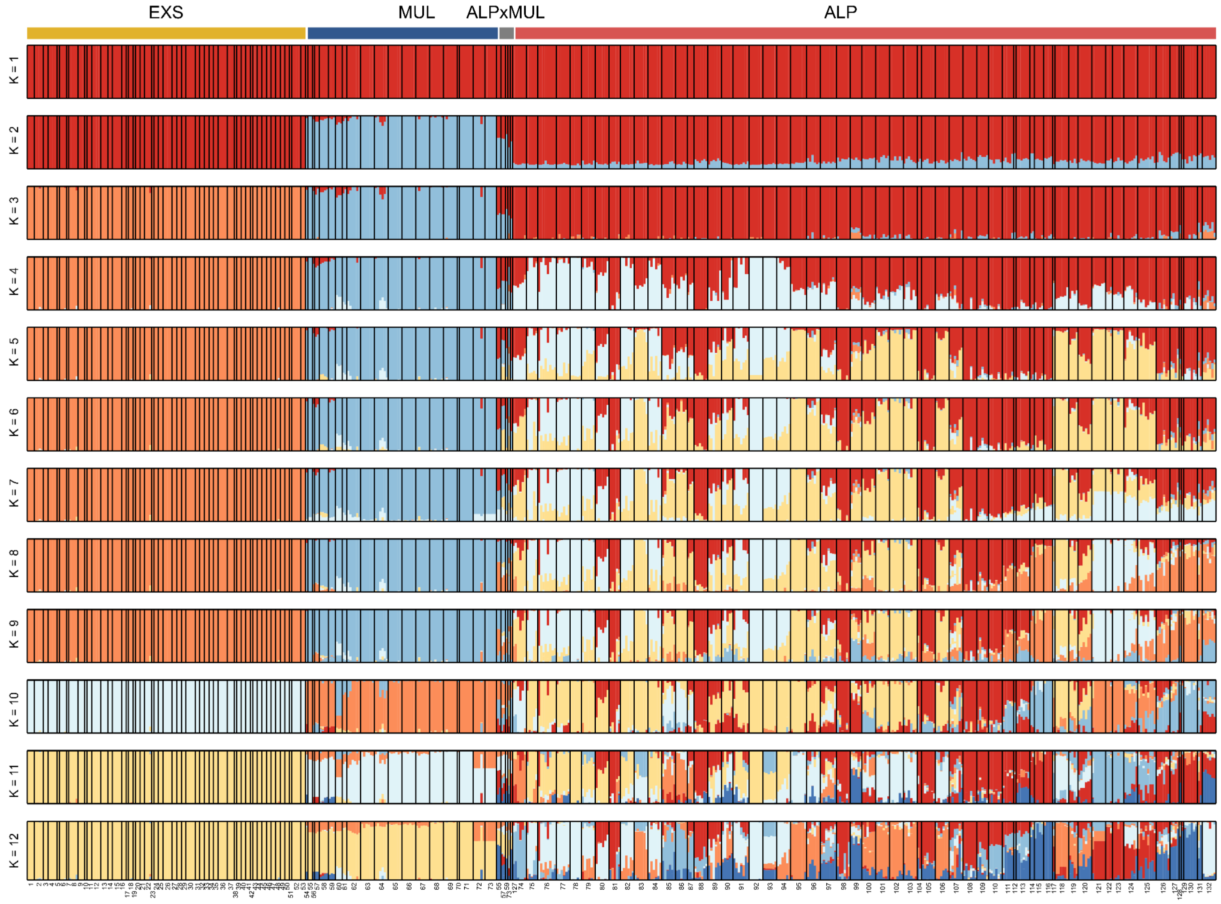

New Phytologist

**Fig. S4** Heatmap of population differentiation (Rho) for *Luzula exspectata* (EXS). Numbers on the sides of the heatmap are population identifiers and dendrograms show clustering based on Euclidean distances of Rho among populations.

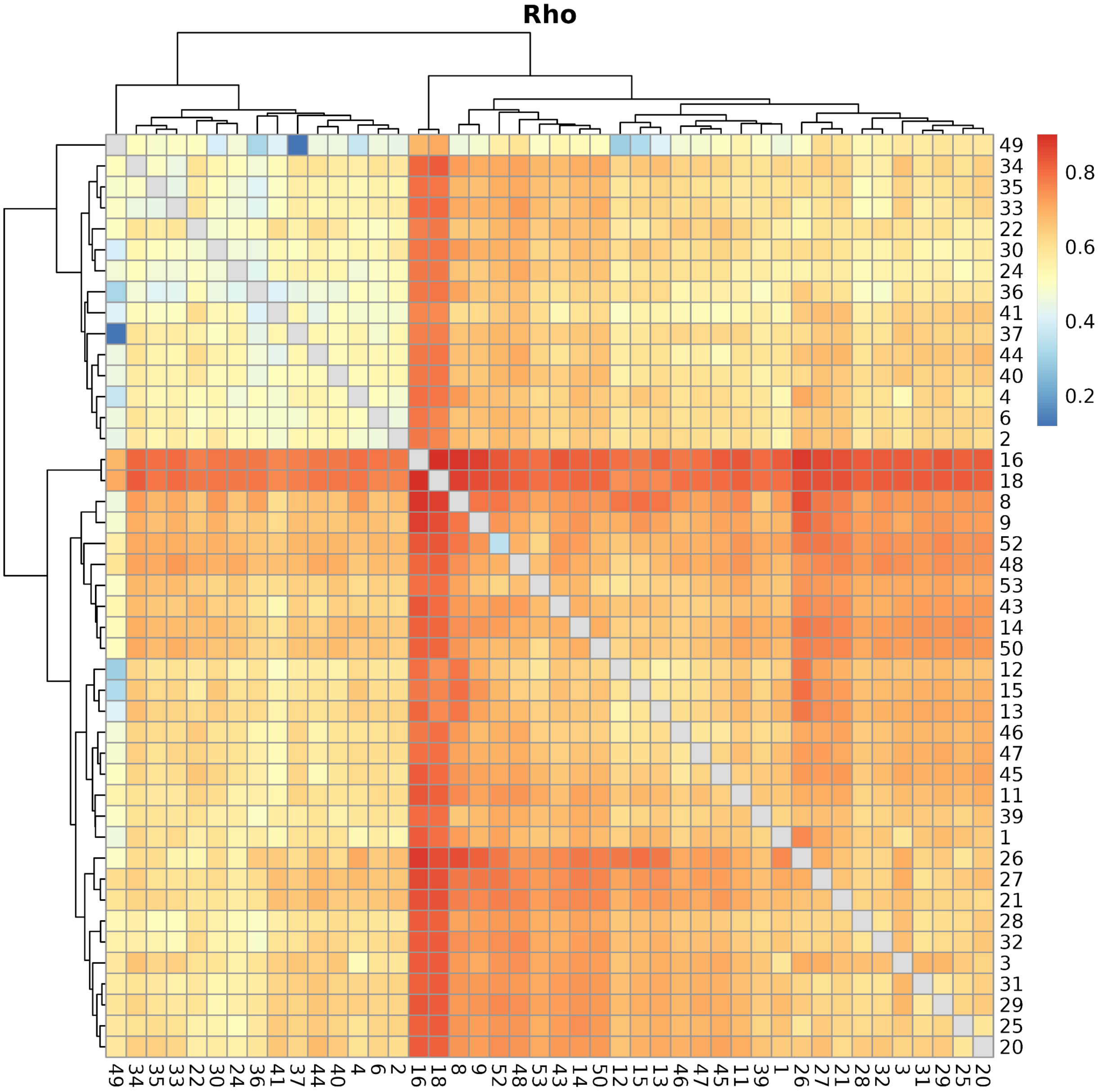

**Fig. S5** Heatmap of population differentiation (Rho) for *Luzula multiflora* (MUL). Numbers on the sides of the heatmap are population identifiers and dendrograms show clustering based on Euclidean distances of Rho among populations.

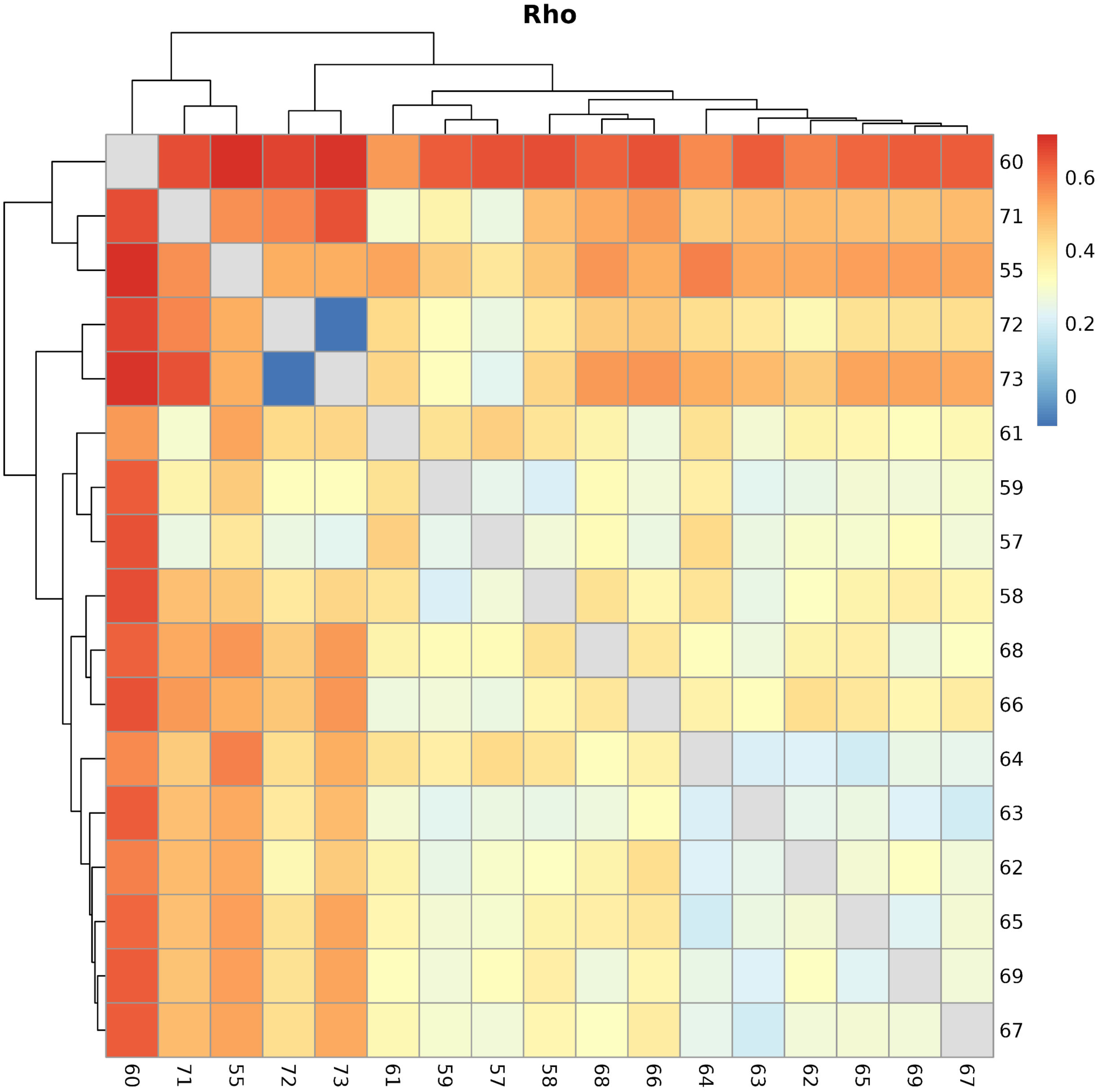

New Phytologist

**Fig. S6** Heatmap of population differentiation (Rho) for *Luzula alpina* (ALP). Numbers on the sides of the heatmap are population identifiers and dendrograms show clustering based on Euclidean distances of Rho among populations.

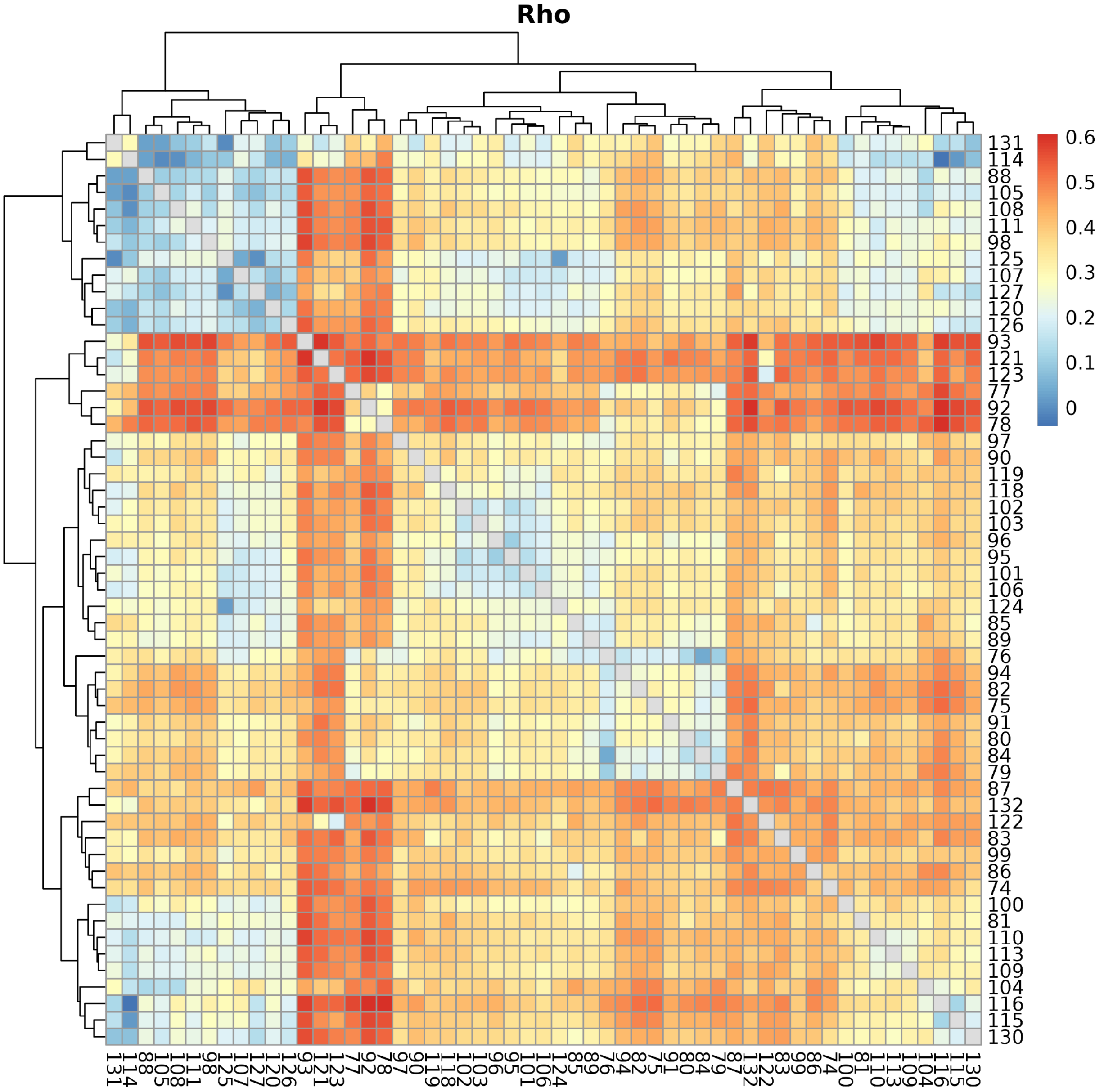

**Fig. S7** Isolation by distance (IBD) for *Luzula exspectata* (EXS; a), *L. multiflora* (MUL; b) and *L. alpina* (ALP; c). Red lines show fitted linear models and grey ribbons are 95% confidence intervals of model predictions. The test statistic (Mantel r) and *P*-value of Mantel tests with 20,000 permutations are shown in the top-left corner.

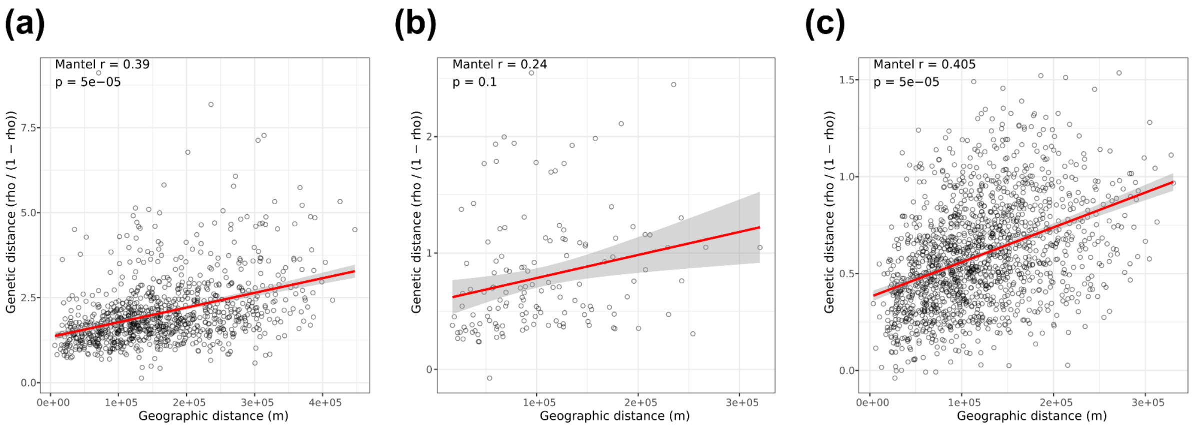


**Fig. S8** Isolation by environment (IBE) for *Luzula exspectata* (EXS; a), *L. multiflora* (MUL; b) and *L. alpina* (ALP; c). Red lines show fitted linear models and grey ribbons are 95% confidence intervals of model predictions. The test statistic (Mantel r) and *P*-value of Mantel tests with 20,000 permutations while accounting for geographic structure (G ~ E | Geo) are shown in the top-left corner.

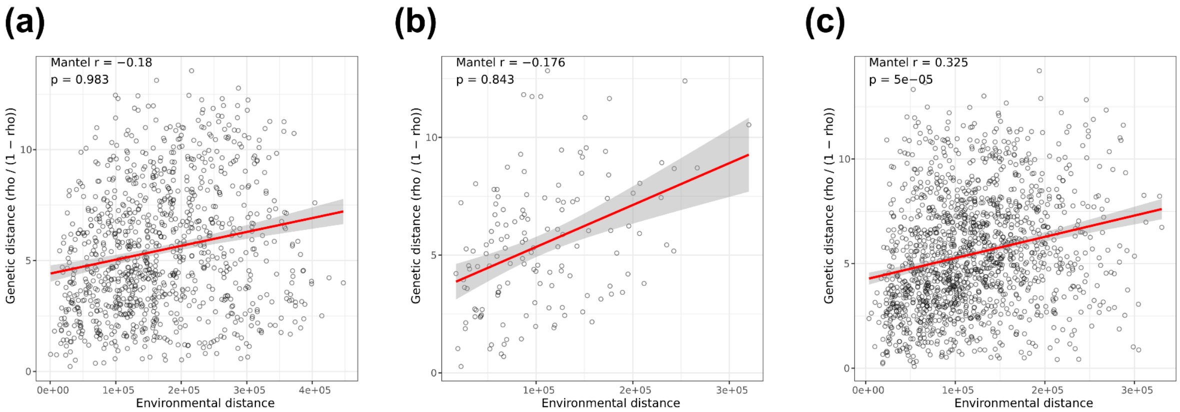

Fig. S9 STRUCTURE results for *Luzula exspectata* (EXS) for $K$ = 1 to 12. Numbers below bar charts are population identifiers.

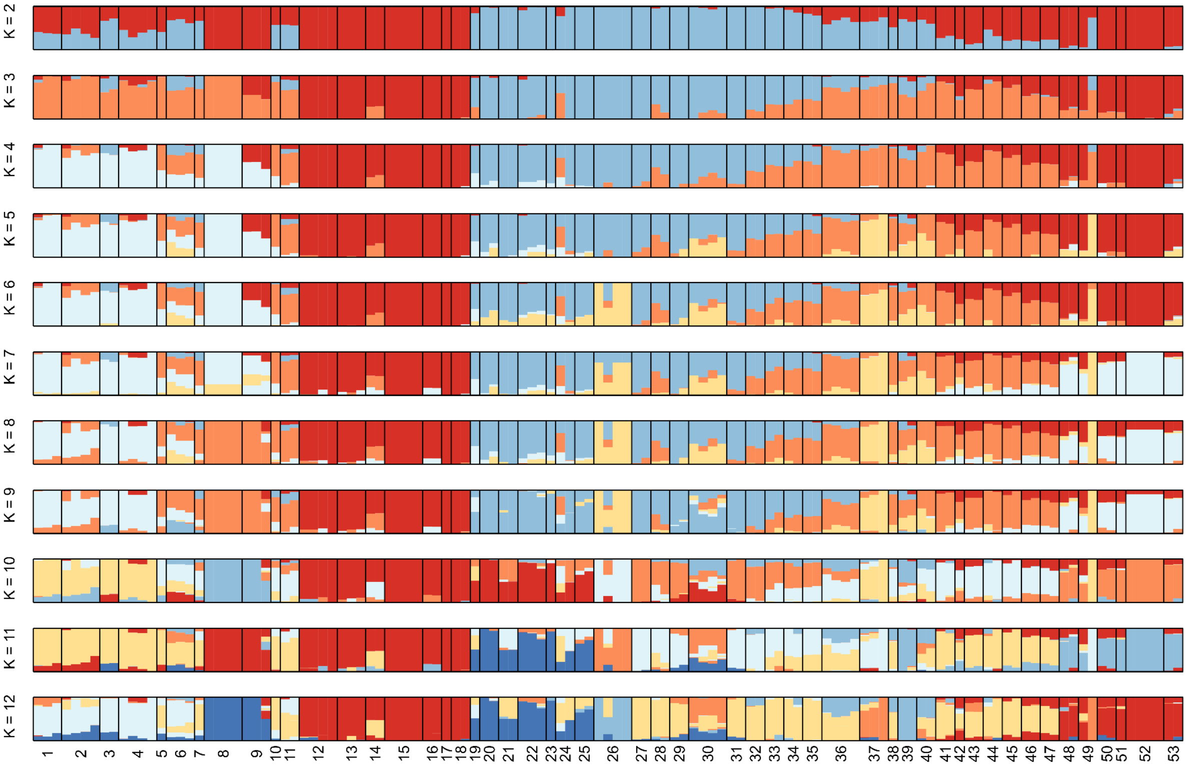

New Phytologist

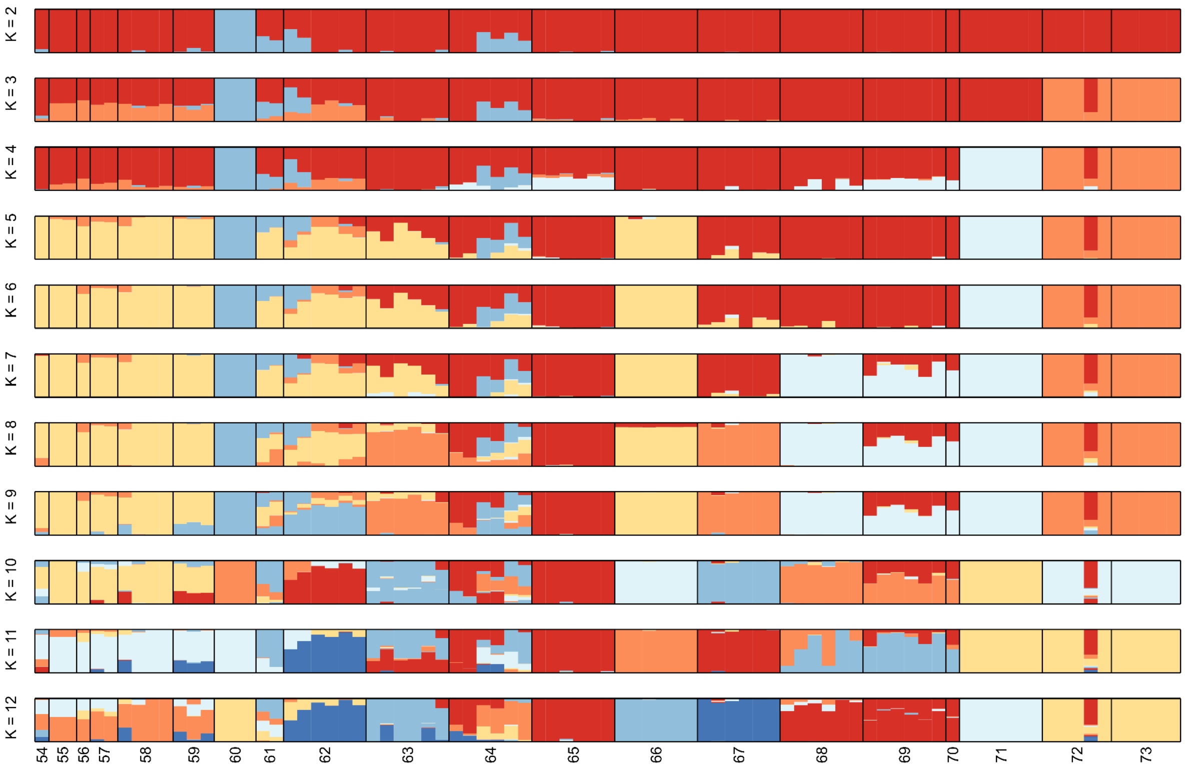


**Fig. S10** STRUCTURE results for *Luzula multiflora* (MUL) for *K* = 1 to 12. Numbers below bar charts are population identifiers.

**Fig. S11** STRUCTURE results for *Luzula alpina* (ALP) for *K* = 1 to 12. Numbers below bar charts are population identifiers.

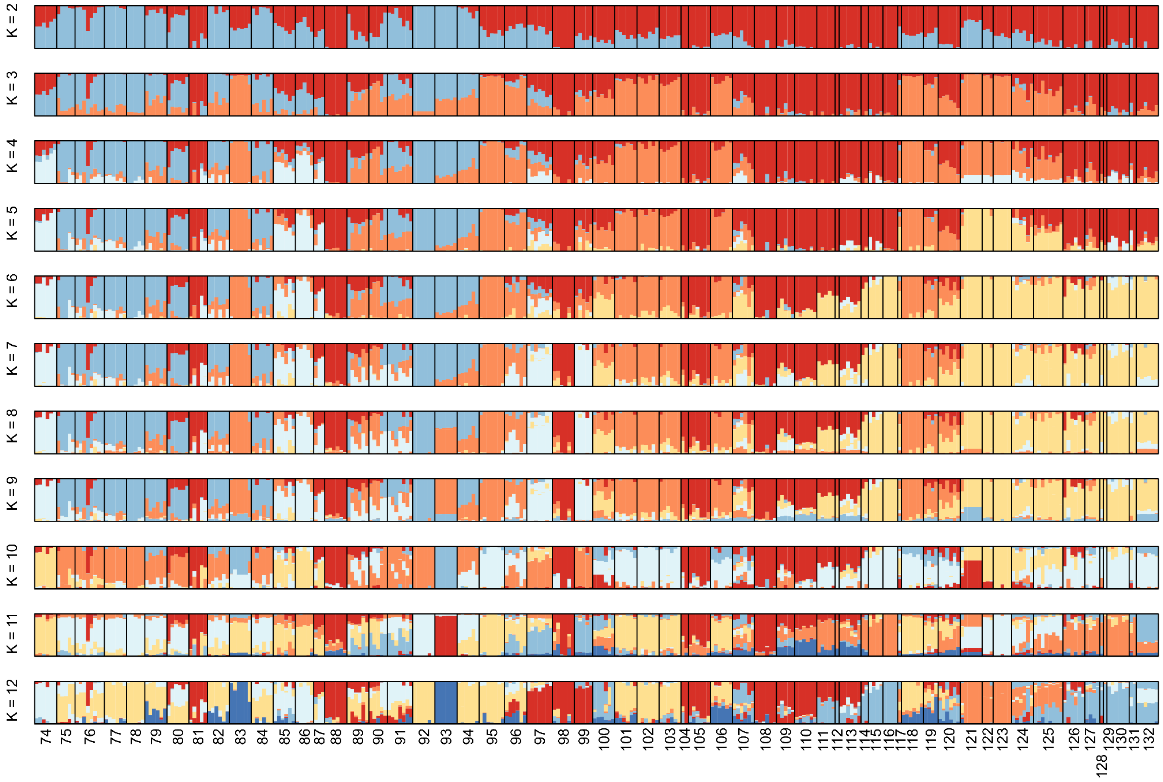

**Fig. S12** Most likely origins of range expansion inferred from the directionality index ψ for either the entire species or major genetic groups defined by STRUCTURE for *Luzula exspectata* (EXS) (a), *L. multiflora* (MUL; b) and *L. alpina* (ALP) (c). Only the full species was tested for MUL as STRUCTURE clusters did not follow clear geographic patterns in this species. Brighter colour shading indicates a higher probability of an area being the origin of range expansion and the most likely origin is highlighted by an X.

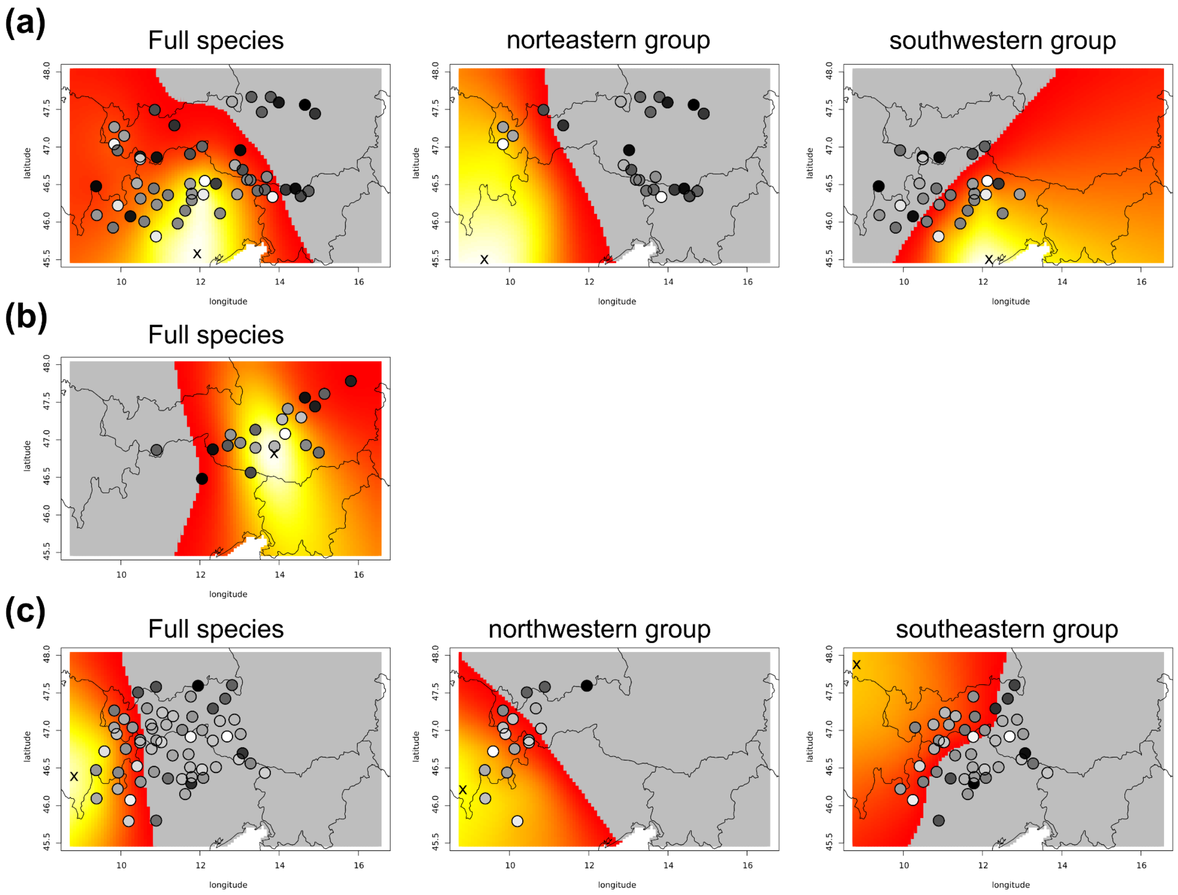

**Fig. S13** Relationship between population-average nucleotide diversity (π) and geographic distance to the inferred origin of range expansion for *Luzula exspectata* (EXS; a), *L. multiflora* (MUL; b) and *L. alpina* (ALP; c). Blue lines show fitted linear models and grey ribbons are 95% confidence intervals of model predictions.

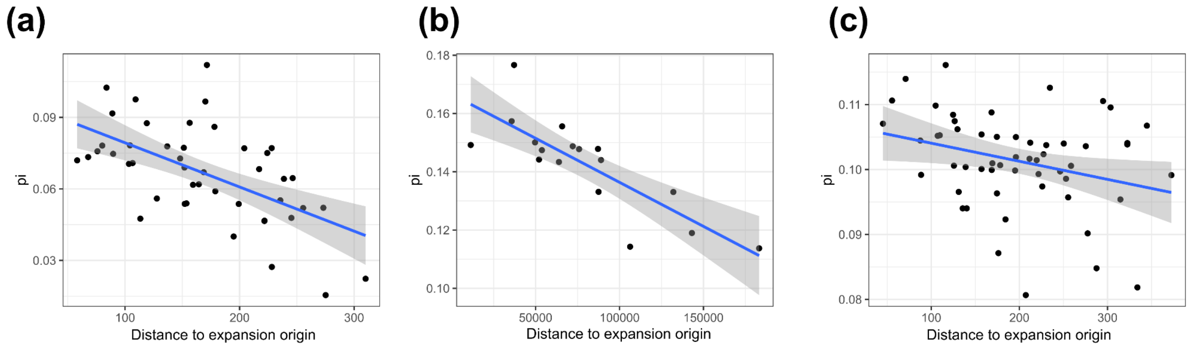


**Fig. S14** Analysis of niches based on coarse-grained ecological data. (a) Pairwise comparison of ecological niches derived from climatic and edaphic ecological data in *ecospat*. Coloured regions show ecological niches for each species along the first two principal components of multivariate niche space while overlapping niches are shown in grey. The background area is shown as a red line and coloured symbols are species occurrences. (b) Variable contributions to the principal component analyses of environmental space.

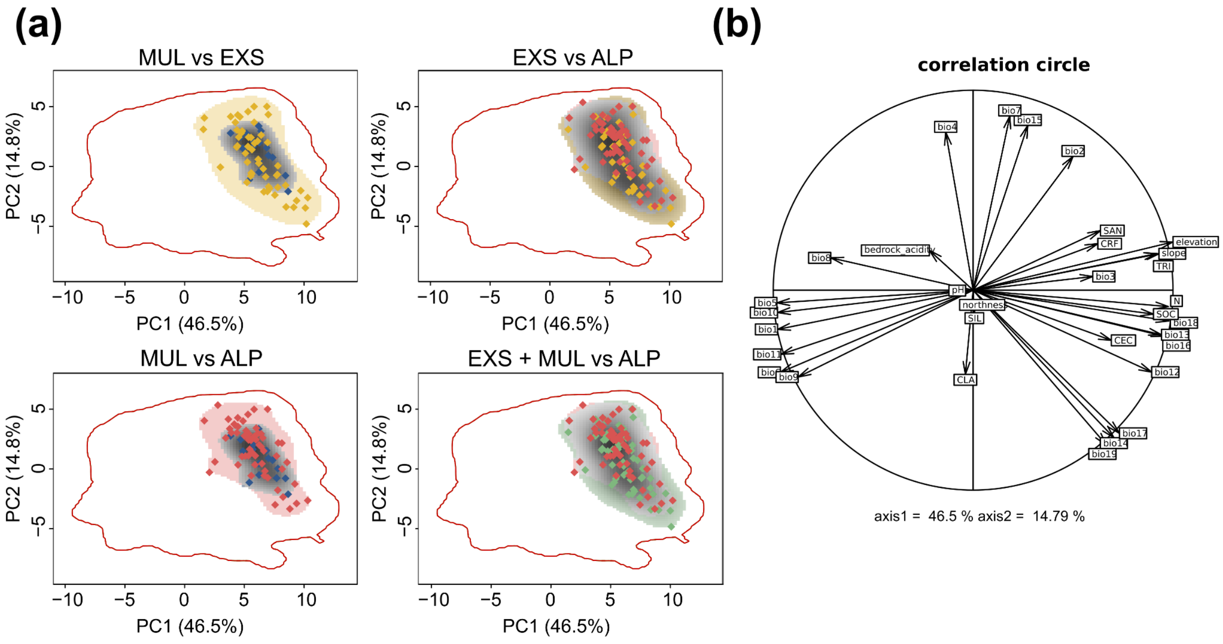

**Fig. S15** Optimised redundancy analysis (RDA) of plant community composition showing the accompanying plant species most strongly differentiating between the three study species. The occurrence of each *Luzula* species shown as colored dots (*Luzula exspectata*, EXS, yellow; *L. multiflora*, MUL, blue; and *L. alpina*, ALP, red) was used as a response variable. The proportion of variation explained by each RDA axis is shown in parentheses.

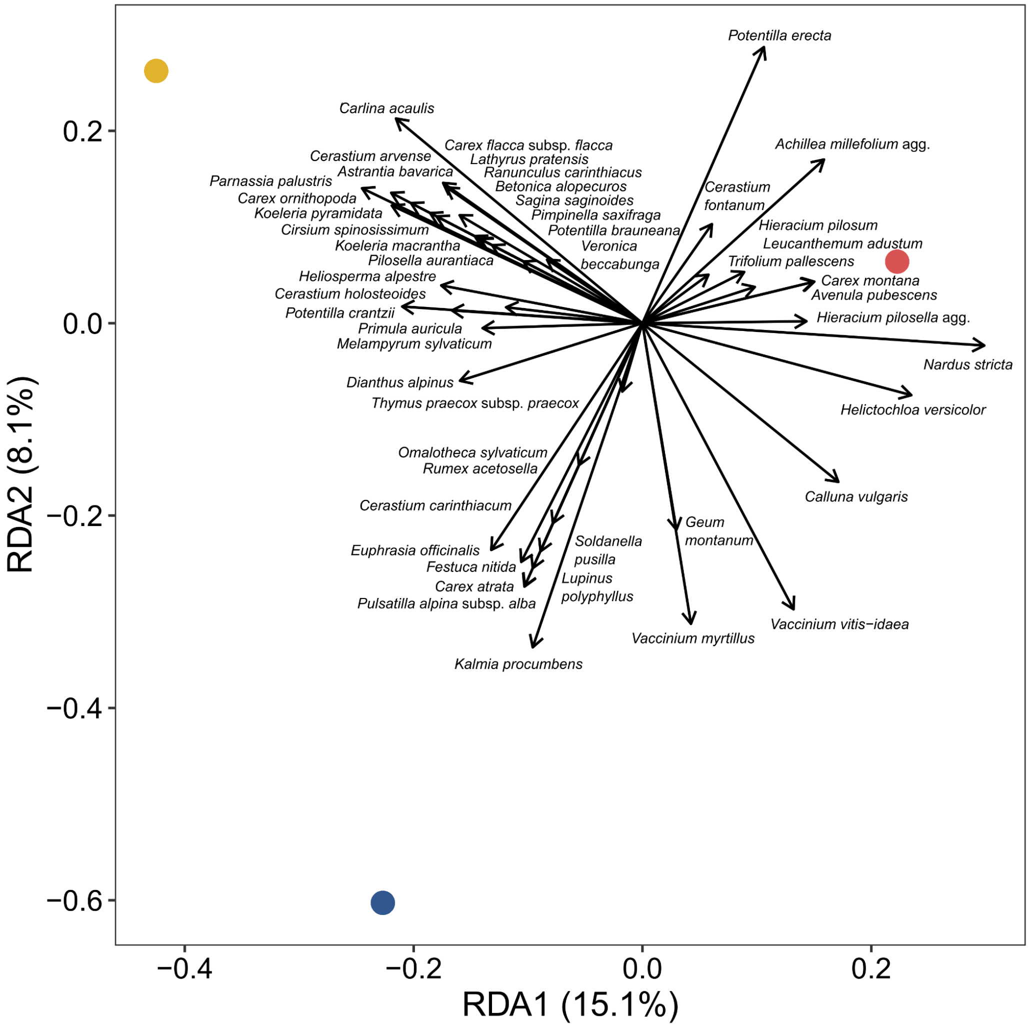

**Fig. S16** Analysis of niches based on fine-grained environmental data. (a) Pairwise comparison of ecological niches computed in *ecospat*. Coloured regions show ecological niches for each species along the first two principal components of multivariate niche space while overlapping niches are shown in grey. The background area is shown as a red line and coloured symbols are species occurrences. (b) Variable contributions to the principal component analyses of environmental space.

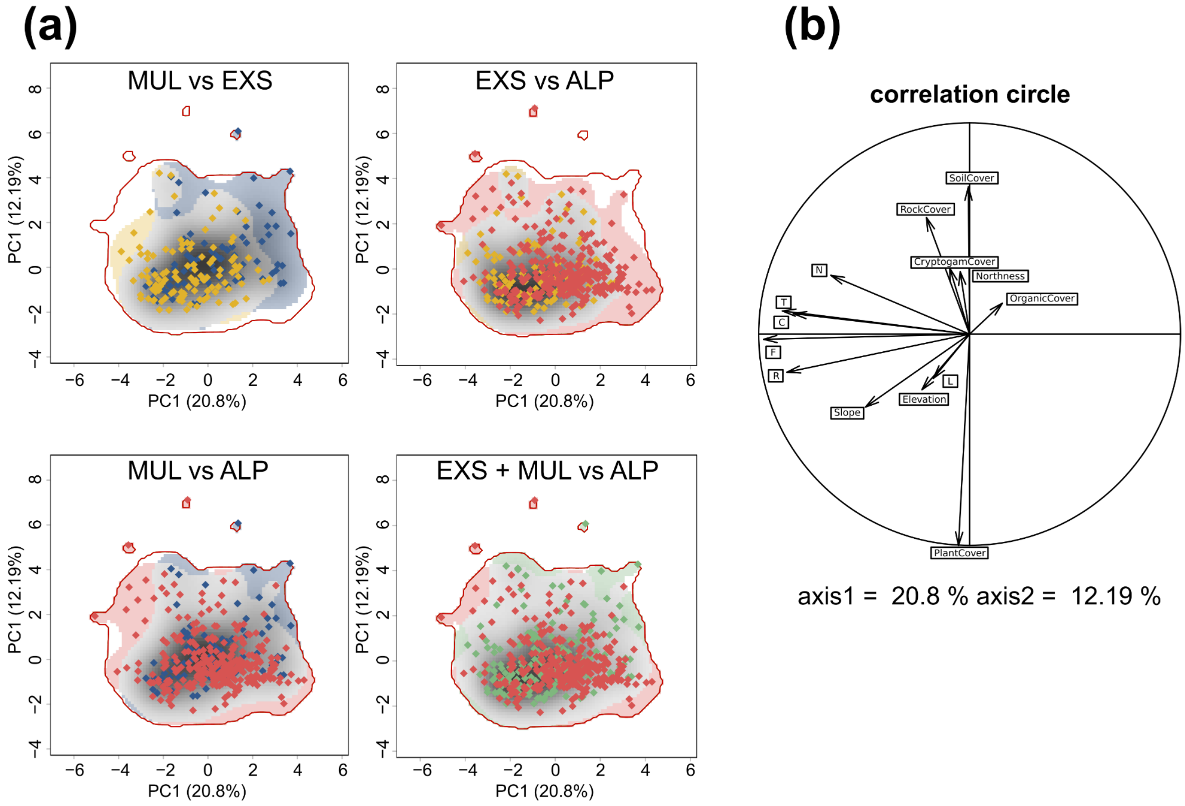

New Phytologist

**Fig. S17** Boxplots of fine-grained environmental data recorded for each individual of *Luzula exspectata* (EXS; yellow), *L. multiflora* (MUL; blue) and *L. alpina* (ALP; red). Community-weighted means of Karrer indicator values were computed from accompanying vascular plant species. Thick horizontal lines are median values, boxes are interquartile ranges (IQR) and whiskers show 1.5 times the IQR. Asterisks above brackets indicate significant mean differences according to Kruskal-Wallis test followed by Dunn's *post hoc* test (ns: not significant, * $P < 0.05$, ** $P < 0.01$, *** $P < 0.001$).

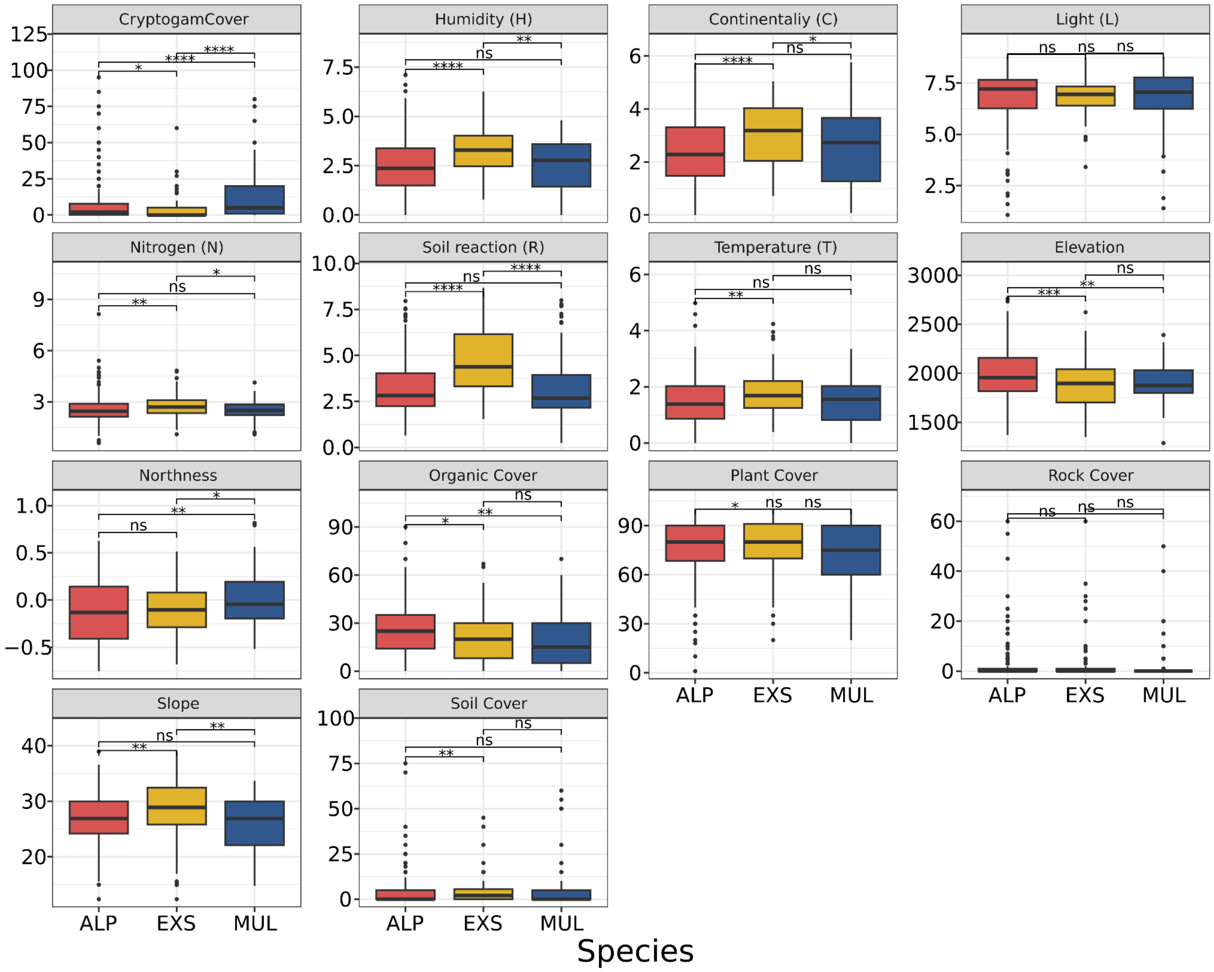

New Phytologist

**Fig. S18** Principal component analysis (PCA) of non-seed morphological characters of *Luzula exspectata* (EXS; yellow), *L. multiflora* (MUL; blue) and *L. alpina* (ALP; red).

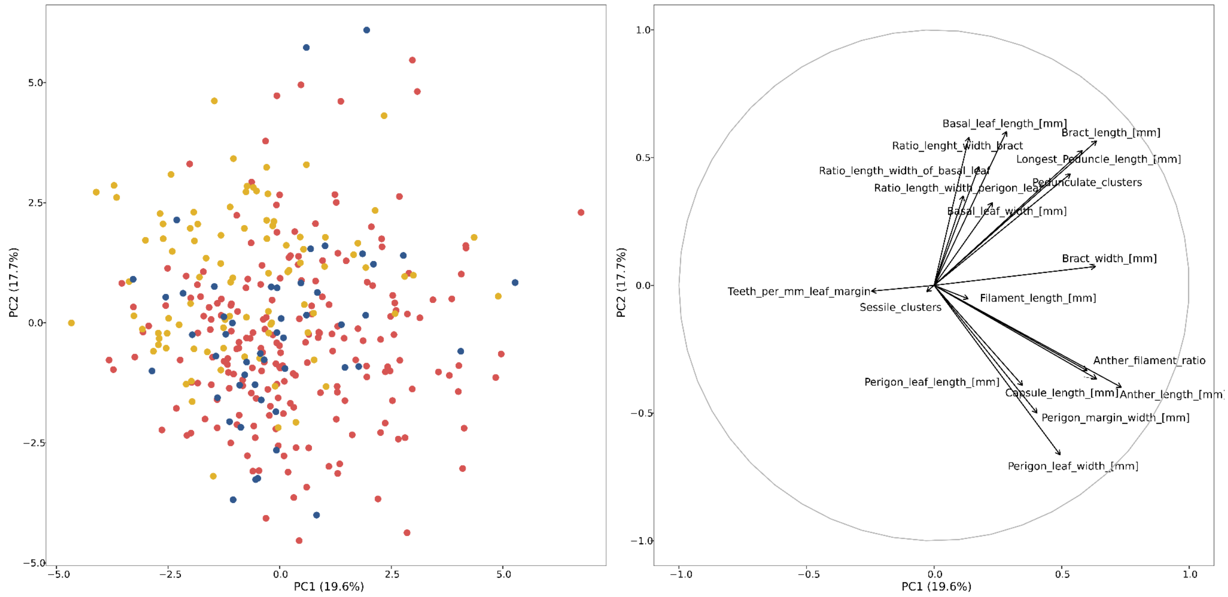

**Fig. S19** Box plots of non-seed morphological characters of *Luzula exspectata* (EXS; yellow), *L. multiflora* (MUL; blue) and *L. alpina* (ALP; red). Thick horizontal lines are median values, boxes are interquartile ranges (IQR) and whiskers show 1.5 times the IQR. Asterisks above brackets indicate significant mean differences according to Kruskal-Wallis test followed by Dunn's *post hoc* test (ns: not significant, * $P < 0.05$, ** $P < 0.01$, *** $P < 0.001$).

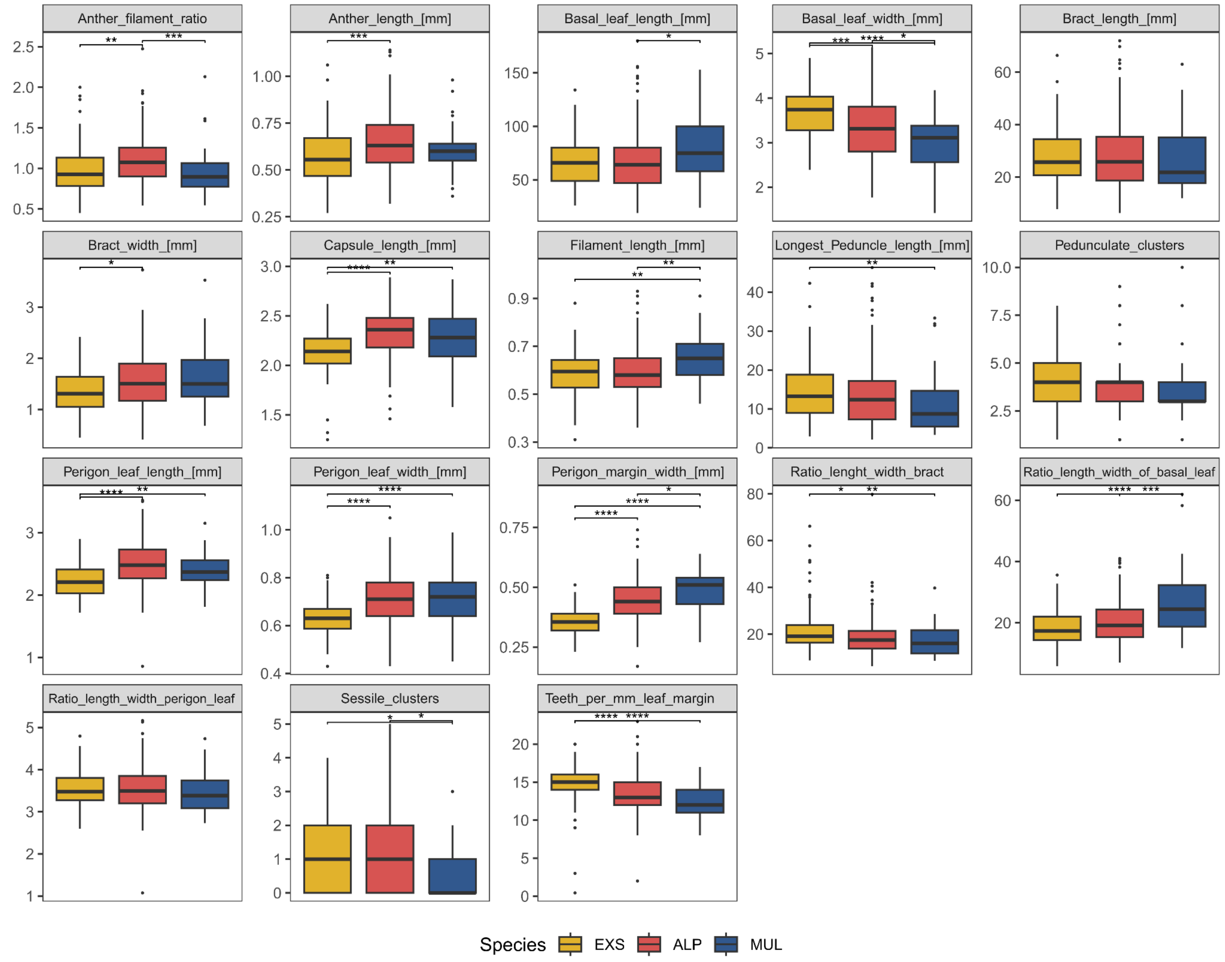

**Fig. S20** Box plots of seed morphological characters of *Luzula exspectata* (EXS; yellow), *L. multiflora* (MUL; blue) and *L. alpina* (ALP; red). Thick horizontal lines are median values, boxes are interquartile ranges (IQR) and whiskers show 1.5 times the IQR. Asterisks above brackets indicate significant mean differences according to Kruskal-Wallis test followed by Dunn's *post hoc* test (ns: not significant, * $P < 0.05$, ** $P < 0.01$, *** $P < 0.001$).

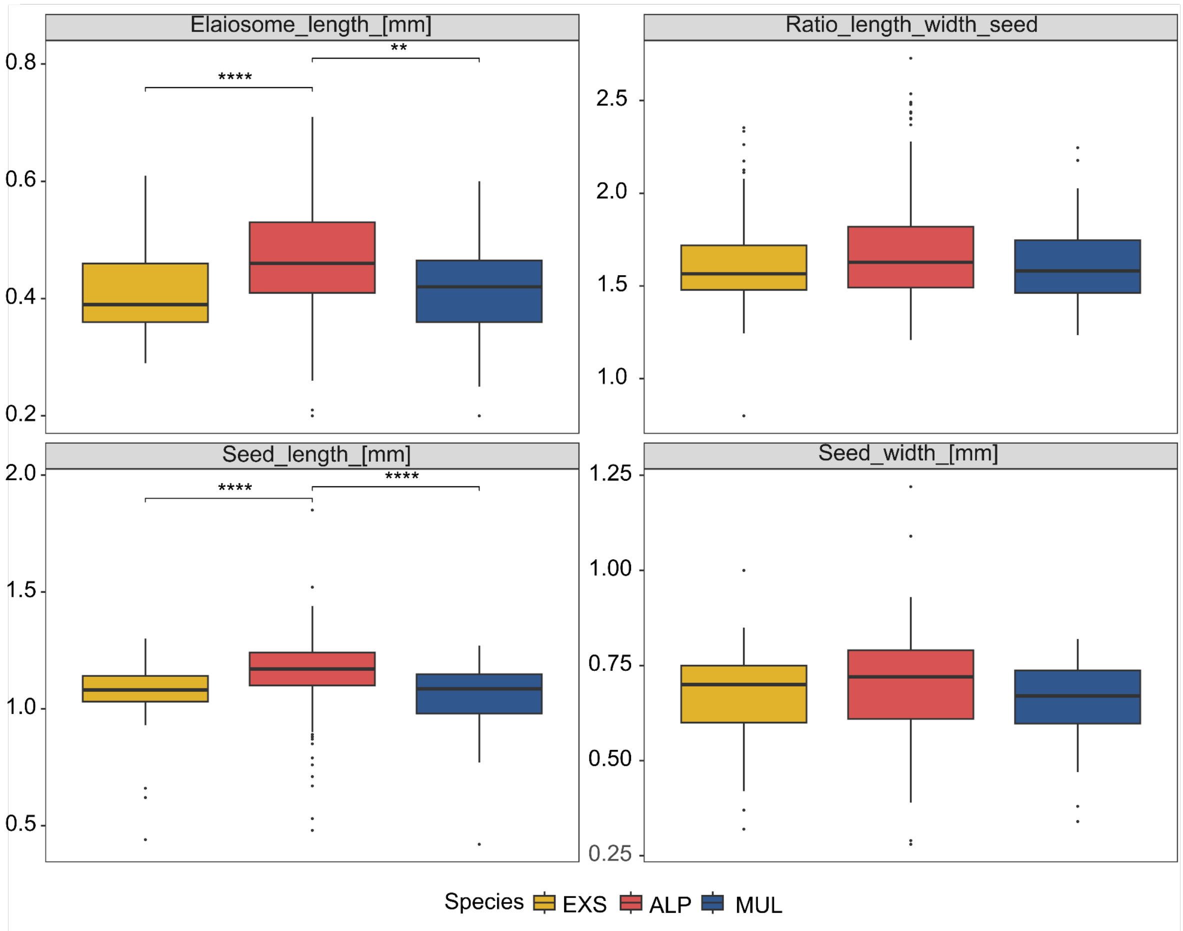

New Phytologist

**Fig. S21** Principal component analysis (PCA) of seed morphological characters of *Luzula exspectata* (EXS; yellow), *L. multiflora* (MUL; blue) and *L. alpina* (ALP; red).

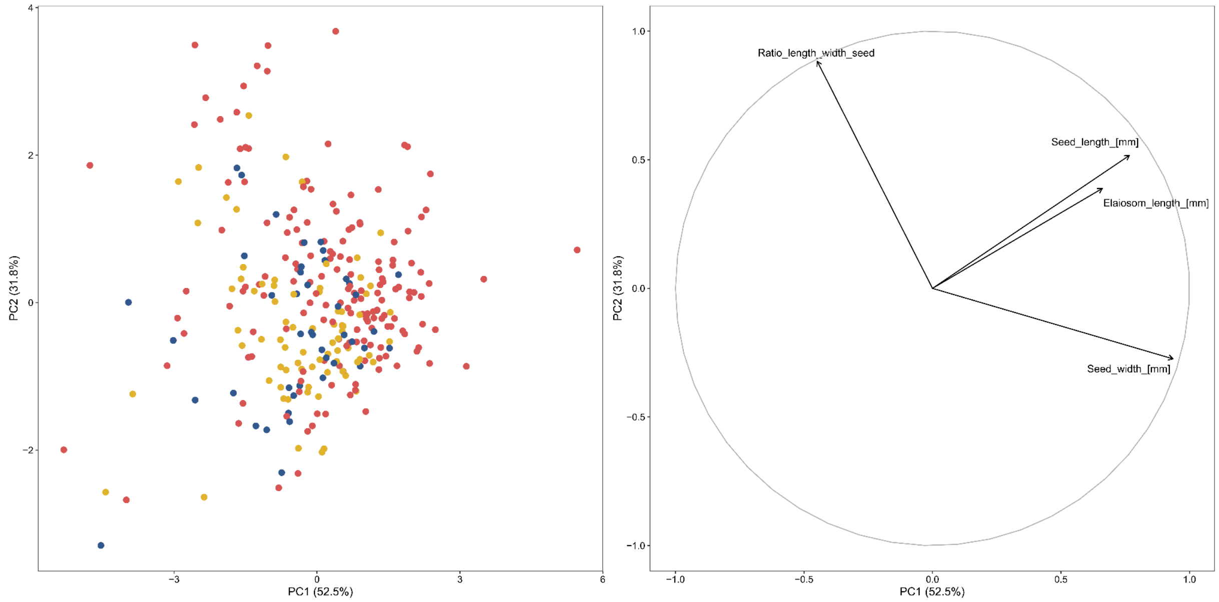


**Fig. S22** Linear discriminant analysis (LDA) of seed morphological characters of *Luzula exspectata* (EXS; yellow), *L. multiflora* (MUL; blue) and *L. alpina* (ALP; red). Ellipses show 95%-quantiles of samples for each species.

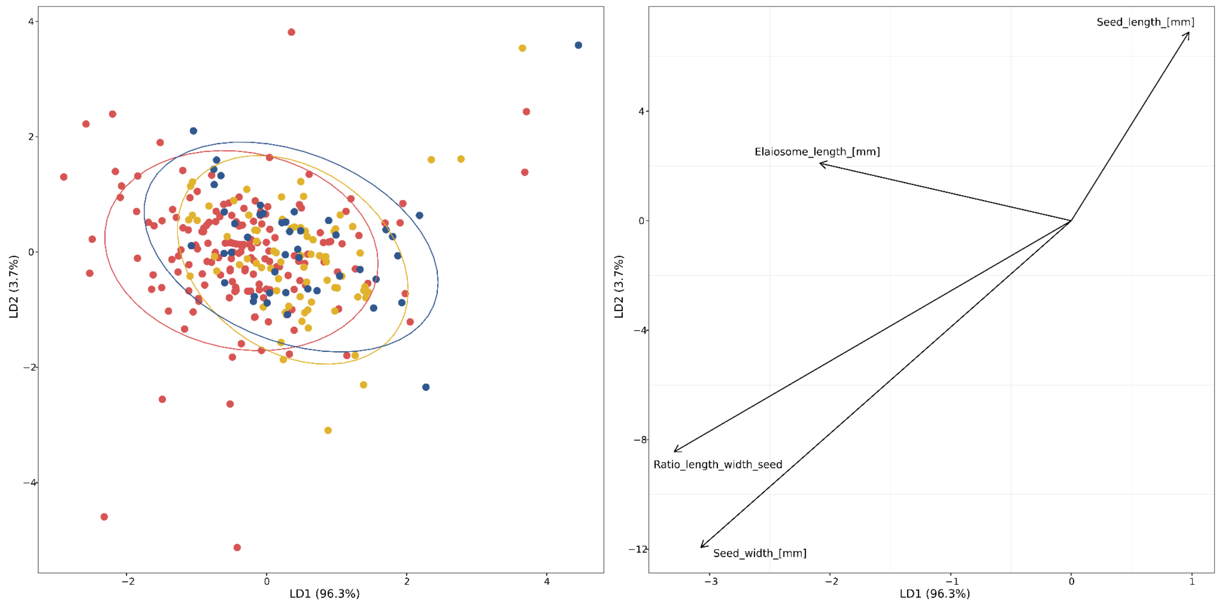

**Table S1** *Luzula* accessions used in this study. This table is available as a separate spreadsheet

**Table S2** Details of filtering applied to ddRADseq data and number of samples and SNPs used for each analysis.

| **Analysis** | **# of SNPs** | **# of samples** | **Filtering** |
|---|---|---|---|
| Baseline VCF for all downstream analyses after quality filtering | 840912 | 517 | GATK hard filtering recommendations "QUAL < 40.0", "QD < 2.0", "MQ < 40.0", "MQRankSum < -12.5", "ReadPosRankSum < -15.0", Filter (and set to no call) genotypes with minDP < 8 and maxDP > 200 |
| NeighbourNet & PCA – all species | 49884 | 517 | ≤ 50% missing data |
| Hybrid index | 2096 | 493 | 1 SNP per RAD locus, only SNPs that differed between EXS and MUL |
| STRUCTURE – all species | 2070 | 517 | ≤ 20% missing data, MAC ≥ 3, 1 SNP per RAD locus |
| STRUCTURE – EXS | 4288 | 121 | ≤ 20% missing data, MAC ≥ 3, 1 SNP per RAD locus |
| STRUCTURE – MUL | 1815 | 83 | ≤ 20% missing data, MAC ≥ 3, 1 SNP per RAD locus |
| STRUCTURE – ALP | 2302 | 306 | ≤ 20% missing data, MAC ≥ 3, 1 SNP per RAD locus |
| Range Expansion – EXS | 2242 | 124 | ≤ 20% missing data, 1 SNP per RAD locus, including outgroup |
| Range Expansion – MUL | 2242 | 86 | ≤ 20% missing data, 1 SNP per RAD locus, including outgroup |
| Range Expansion – ALP | 2242 | 309 | ≤ 20% missing data, 1 SNP per RAD locus, including outgroup |
| Population statistics – EXS | 5419 | 121 | ≤ 20% missing data, 1 SNP per RAD locus, min. 2 individuals per pop |
| Population statistics – MUL | 1965 | 83 | ≤ 20% missing data, 1 SNP per RAD locus, min. 2 individuals per pop |
| Population statistics – ALP | 2440 | 306 | ≤ 20% missing data, 1 SNP per RAD locus, min. 2 individuals per pop |

New Phytologist

**Table S3** Climatic and edaphic variables used for environmental niche modelling (ENM) and niche analysis in *ecospat*.

| ENM | Niche analysis | Abbreviation | Definition |
|---|---|---|---|
| 1 | 1 | bio1 | Annual Mean Temperature [°C] |
| | 1 | bio2 | Mean Diurnal Range [°C] |
| | 1 | bio3 | Isothermality |
| | 1 | bio4 | Temperature Seasonality [standard deviation] |
| | 1 | bio5 | Max Temperature of Warmest Month [°C] |
| | 1 | bio6 | Min Temperature of Coldest Month [°C] |
| | 1 | bio7 | Temperature Annual Range [°C] |
| 1 | 1 | bio8 | Mean Temperature of Wettest Quarter [°C] |
| | 1 | bio9 | Mean Temperature of Driest Quarter [°C] |
| | 1 | bio10 | Mean Temperature of Warmest Quarter [°C] |
| | 1 | bio11 | Mean Temperature of Coldest Quarter [°C] |
| 1 | 1 | bio12 | Annual Precipitation [mm/year] |
| | 1 | bio13 | Precipitation of Wettest Month [mm/month] |
| | 1 | bio14 | Precipitation of Driest Month [mm/month] |
| 1 | 1 | bio15 | Precipitation Seasonality [coefficient of variation] |
| | 1 | bio16 | Precipitation of Wettest Quarter [mm/quarter] |
| | 1 | bio17 | Precipitation of Driest Quarter [mm/quarter] |
| | 1 | bio18 | Precipitation of Warmest Quarter [mm/quarter] |
| | 1 | bio19 | Precipitation of Coldest Quarter [mm/quarter] |
| 1 | 1 | bedrock_acidity | Bedrock acidity computed from Geo-Lim |
| | 1 | CEC | Cation exchange capacity at pH7 at 5-15cm soil depth [mmol(c)/kg] |
| | 1 | CLA | Clay content at 5-15cm soil depth [g/kg] |
| | 1 | CRF | Coarse fragments at 5-15cm soil depth [$cm^3/dm^3$] |
| | 1 | N | Nitrogen at 5-15cm soil depth [cg/kg] |
| | 1 | pH | pH in water at 5-15cm soil depth [pH*10] |
| | 1 | SAN | Sand content at 5-15 cm soil depth [g/kg] |
| | 1 | SIL | Silt content at 5-15 cm soil depth [g/kg] |
| | 1 | SOC | Soil organic carbon at 5-15 cm soil depth [dg/kg] |

**Table S4** Vegetation relevé data. This table is available as a separate spreadsheet.

**Table S5** Fine-grained ecological data. This table is available as a separate spreadsheet.

**Table S6** Taxa with no Karrer ecological indicator values (EIVs) available. Adapted EIVs were computed for four species by rescaling Landolt EIVs to the range of Karrer EIVs (8 records, 0.1% of data). Three species groups without EIV were excluded from niche analyses (170 records, 2.8% data). L: light availability; T: temperature; C: continentality; H: humidity; R: soil reaction; N: nitrogen availability.

| **Taxon** | **adapted Landolt EIVs** | | | | | | | **no EIVs available** |
|---|---|---|---|---|---|---|---|---|
| | **L** | **T** | **C** | **H** | **R** | **N** | **# samples** | **# samples** |
| *Hypericum richeri* | 5 | 5 | 4 | 5 | 9 | 5.4 | 1 | |
| *Gentianella ramosa* | 7 | 3 | 7 | 5 | 3.6 | 3.6 | 1 | |
| *Primula glaucescens* | 7 | 4 | 5 | 6 | 7.2 | 3.6 | 1 | |
| *Astrantia minor* | 5 | 5 | 5 | 5 | 3.6 | 3.6 | 5 | |
| *Alchemilla alpina* agg. | | | | | | | | 19 |
| *Alchemilla vulgaris* agg. | | | | | | | | 97 |
| *Hieracium pilosella* agg. | | | | | | | | 54 |
| Total | | | | | | | 8 (0.1%) | 170 (2.8%) |

**Table S7** Morphometric data. This table is available as a separate spreadsheet.

**Table S8** Results of Mantel tests with 20,000 permutations to test for the effects of isolation by distance (IBD) and isolation by environment (IBE). IBD was inferred from correlations of genetic distances (rho/(rho-1)) and Euclidean geographic distances (G ~ Geo). IBE was calculated from the correlation of genetic distances and Euclidean distances from principal components of environmental space while accounting for spatial structure (G ~ E | Geo). Mantel r statistics and *P*-values are reported for all tests. *P*-values < 0.05 are indicated with bold font.

| | **G ~ Geo (IBD)** | | **G ~ E** | | **G ~ E \| Geo (IBE)** | |
|---|---|---|---|---|---|---|
| **Species** | **r** | ***P*-value** | **r** | ***P*-value** | **r** | ***P*-value** |
| EXS | 0.3904 | **9.999e-05** | -0.08187 | 0.81932 | -0.2245 | 0.9956 |
| MUL | 0.2403 | 0.09679 | 0.261 | 0.075092 | 0.1671 | 0.16218 |
| ALP | 0.4054 | **9.999e-05** | 0.4227 | **9.999e-05** | 0.3195 | **9.999e-05** |

**Table S9** Results of multivariate redundancy analysis associating population-level allele frequencies with geographic coordinates and/or coordinates along the first two principal components of environmental space. Significance was assessed with 20,000 permutations. Isolation by distance (IBD) was inferred from correlations of allele frequencies and geographic coordinates (G ~ Geo). IBE was calculated from the correlation of allele frequencies and coordinates in environmental space while accounting for spatial structure (G ~ E | Geo). F statistics and *P*-values are reported for all tests. *P*-values < 0.05 are indicated with bold font.

| | **G ~ Geo (IBD)** | | **G ~ E** | | **G ~ E \| Geo (IBE)** | |
|---|---|---|---|---|---|---|
| **Species** | **F-statistic** | ***P*-value** | **F-statistic** | ***P*-value** | **F-statistic** | ***P*-value** |
| EXS | 2.748 | **0.0000** | 1.957 | **0.0001** | 1.255 | **0.0435** |
| MUL | 1.941 | **0.0001** | 1.456 | **0.0176** | 1.244 | 0.1340 |
| ALP | 4.703 | **0.0000** | 1.744 | **0.0015** | 1.719 | **0.0002** |

**Table S10** Inferred approximate geographic origins and results of the test for geographic expansion for the entire species and the two major STRUCTURE clusters for *L. alpina* (ALP) and *L. exspectata* (EXS). *q* is the strength of the directionality index, R100 is the estimated decrease in genomic diversity over 100 km, and $R^2$ and *P* are the correlation coefficient and *P*-value for the most likely geographic origin, respectively. *P*-values < 0.05 are indicated with bold font.

| Species | Region | Origin | q | R100 | $R^2$ | *P* |
|---|---|---|---|---|---|---|
| EXS | full | 45.5758° N, 11.9111° E | 0.0001 | 0.9806 | 0.0522 | **1.24e-18** |
| | northeast | 45.5000° N, 9.3444° E | 0.0002 | 0.9633 | 0.1637 | **1.86e-14** |
| | southwest | 45.5000° N, 12.1444° E | 0.0002 | 0.9474 | 0.1669 | **7.43e-17** |
| MUL | full | 46.8131° N, 13.8556° E | 0.0004 | 0.9217 | 0.5617 | **9.93e-36** |
| ALP | full | 46.3838° N, 8.8000 ° E | 0.0002 | 0.9615 | 0.1330 | **3.49e-55** |
| | northwest | 46.2071° N, 8.8000 ° E | 0.0005 | 0.9162 | 0.4961 | **6.67e-22** |
| | southeast | 47.8737° N, 8.8000 ° E | 0.0001 | 0.9764 | 0.0399 | **1.98e-09** |

**Table S11** Summary of linear models to test for the relationship between population-average nucleotide diversity (π) and geographic distance to the inferred origin of range expansion. *P*-values < 0.05 are indicated with bold font.

| Species | Model | Term | Estimate | Std. Error | t-value | *P*-value | df | adj. $R^2$ | F-statistic |
|---|---|---|---|---|---|---|---|---|---|
| EXS | pi ~ distance | intercept | 9.787e-02 | 7.075e-03 | 13.832 | **< 2e-16** | 43 | 0.3251 | 22.2 |
| | | distance | -1.853e-04 | 3.933e-05 | -4.712 | **2.59e-05** | | | |
| MUL | pi ~ distance | intercept | 1.666e-01 | 5.079e-03 | 32.801 | **2.22e-15** | 15 | 0.631 | 28.35 |
| | | distance | -3.022e-07 | 5.675e-08 | -5.325 | **8.49e-05** | | | |
| ALP | pi ~ distance | intercept | 1.068e-01 | 2.596e-03 | 41.155 | **< 2e-16** | 53 | 0.07181 | 5.177 |
| | | distance | -2.787e-05 | 1.225e-05 | -2.275 | **0.027** | | | |

**Table S12** Metrics of niche divergence computed from 32 climatic, topographic, lithologic and edaphic variables from the coarse-grained dataset for all pairwise comparisons among study species and between *Luzula alpina* (ALP) and the combined niche of *L. exspectata* (EXS) and *L. multiflora* (MUL). D: Schoener's *D*, I: Hellinger's *I*, $P_{Deq}$: *P*-value of the niche equivalency test for *D*, $P_{Ieq}$: *P*-value of the niche equivalency test for *I*, $P_{Dsim}$: *P*-value of the niche similarity test for *D*, $P_{Isim}$: *P*-value of the niche similarity test for *I*, $B_{PC1}$: niche breadth along PC1 (median ± standard deviation), $B_{PC2}$: niche breadth along PC2, S: niche stability, U: niche unfilling, E: niche expansion. Significant *P*-values < 0.05 are highlighted in bold. Niche breadth is given for the species indicated.

| Comparison | D | I | $P_{Deq}$ | $P_{Ieq}$ | $P_{Dsim}$ | $P_{Isim}$ | $B_{PC1}$ | $B_{PC2}$ | S | U | E |
|---|---|---|---|---|---|---|---|---|---|---|---|
| MUL – EXS | 0.54 | 0.73 | **0.0110** | **0.0150** | **0.0500** | **0.0480** | EXS 2.07±0.13 | EXS 2.54±0.12 | 0.54 | 0.00 | 0.46 |
| EXS – ALP | 0.75 | 0.91 | 0.2927 | 0.3816 | **0.0460** | **0.0340** | ALP 1.83±0.11 | ALP 2.04±0.14 | 0.99 | 0.13 | 0.01 |
| MUL – ALP | 0.63 | 0.79 | 0.0769 | 0.0949 | **0.0280** | **0.0230** | MUL 1.37±0.07 | MUL 1.47±0.07 | 0.66 | 0.00 | 0.34 |
| EXS+MUL – ALP | 0.78 | 0.93 | 0.3037 | 0.4565 | **0.0410** | **0.0330** | 1.95±0.13 | 2.34±0.13 | 0.99 | 0.10 | 0.00 |

**Table S13** ANOVA results for the optimised model, axes and explanatory variables of the redundancy analysis (RDA) of macroenvironmental variables.

| **RDA ANOVA model** | **Df** | **Variance** | **F value** | ***P*-value** |
|---|---|---|---|---|
| Model | 4 | 0.135 | 8.852 | 0.001 |
| Residual | 127 | 0.485 | | |
| **RDA ANOVA axes** | **Df** | **Variance** | **F value** | ***P*-value** |
| RDA1 | 1 | 0.077 | 20.376 | 0.001 |
| RDA2 | 1 | 0.059 | 15.590 | 0.002 |
| Residual | 129 | 0.485 | | |
| **RDA ANOVA explanatory variables** | **Df** | **Variance** | **F value** | ***P*-value** |
| N | 1 | 0.046 | 12.115 | 0.001 |
| bedrock_acidity | 1 | 0.040 | 10.453 | 0.001 |
| bio19 | 1 | 0.025 | 6.519 | 0.010 |
| pH | 1 | 0.024 | 6.320 | 0.003 |
| Residual | 127 | 0.485 | | |

**Table S14** ANOVA results for the optimised model, axes and explanatory variables of the redundancy analysis (RDA) of accompanying plant species.

| **RDA ANOVA model** | **Df** | **Variance** | **F value** | ***P*-value** |
|---|---|---|---|---|
| Model | 49 | 0.231 | 6.655 | 0.001 |
| Residual | 455 | 0.323 | | |
| **RDA ANOVA axes** | **Df** | **Variance** | **F value** | ***P*-value** |
| RDA1 | 1 | 0.151 | 234.23 | 0.001 |
| RDA2 | 1 | 0.081 | 125.56 | 0.001 |
| Residual | 502 | 0.323 | | |
| **RDA ANOVA explanatory variables** | **Df** | **Variance** | **F value** | ***P*-value** |
| *Nardus stricta* | 1 | 0.013 | 19.012 | 0.001 |

| | | | | |
|---|---|---|---|---|
| *Kalmia procumbens* | 1 | 0.010 | 14.198 | 0.001 |
| *Carlina acaulis* | 1 | 0.010 | 13.739 | 0.001 |
| *Parnassia palustris* | 1 | 0.008 | 11.906 | 0.001 |
| *Astrantia bavarica* | 1 | 0.008 | 11.231 | 0.001 |
| *Euphrasia officinalis* | 1 | 0.008 | 10.690 | 0.001 |
| *Carex atrata* | 1 | 0.008 | 10.997 | 0.001 |
| *Festuca nitida* | 1 | 0.007 | 10.506 | 0.001 |
| *Vaccinium myrtillus* | 1 | 0.007 | 9.807 | 0.002 |
| *Soldanella pusilla* | 1 | 0.007 | 9.944 | 0.002 |
| *Helictochloa versicolor* | 1 | 0.007 | 10.354 | 0.001 |
| *Achillea millefolium* agg. | 1 | 0.006 | 9.054 | 0.001 |
| *Potentilla crantzii* | 1 | 0.006 | 7.934 | 0.001 |
| *Lathyrus pratensis* | 1 | 0.006 | 8.856 | 0.001 |
| *Carex flacca* subsp. *flacca* | 1 | 0.006 | 7.984 | 0.003 |
| *Pulsatilla alpina* subsp. *alba* | 1 | 0.006 | 7.786 | 0.001 |
| *Rumex acetosella* | 1 | 0.003 | 4.443 | 0.001 |
| *Lupinus polyphyllus* | 1 | 0.006 | 8.152 | 0.001 |
| *Ranunculus carinthiacus* | 1 | 0.005 | 7.045 | 0.001 |
| *Geum montanum* | 1 | 0.005 | 7.525 | 0.001 |
| *Hieracium pilosella* agg. | 1 | 0.005 | 7.401 | 0.003 |
| *Cirsium spinosissimum* | 1 | 0.004 | 5.275 | 0.001 |
| *Pilosella aurantiaca* | 1 | 0.003 | 4.916 | 0.001 |
| *Dianthus alpinus* | 1 | 0.003 | 4.903 | 0.006 |
| *Koeleria macrantha* | 1 | 0.004 | 5.733 | 0.001 |
| *Calluna vulgaris* | 1 | 0.004 | 5.184 | 0.011 |
| *Carex montana* | 1 | 0.003 | 4.807 | 0.014 |
| *Potentilla erecta* | 1 | 0.003 | 4.698 | 0.005 |
| *Avenula pubescens* | 1 | 0.003 | 3.891 | 0.026 |
| *Heliosperma alpestre* | 1 | 0.003 | 4.177 | 0.012 |
| *Trifolium pallescens* | 1 | 0.003 | 4.560 | 0.016 |
| *Cerastium arvense* | 1 | 0.003 | 4.825 | 0.012 |
| *Carex ornithopoda* | 1 | 0.002 | 3.362 | 0.046 |
| *Hieracium pilosum* | 1 | 0.003 | 4.186 | 0.016 |
| *Vaccinium vitis-idaea* | 1 | 0.003 | 4.339 | 0.017 |

| | | | | |
|---|---|---|---|---|
| *Leucanthemum adustum* | 1 | 0.003 | 4.794 | 0.007 |
| *Cerastium fontanum* | 1 | 0.003 | 4.312 | 0.018 |
| *Potentilla brauneana* | 1 | 0.004 | 4.945 | 0.001 |
| *Cerastium carinthiacum* | 1 | 0.003 | 3.763 | 0.030 |
| *Betonica alopecuros* | 1 | 0.003 | 3.814 | 0.016 |
| *Melampyrum sylvaticum* | 1 | 0.002 | 3.363 | 0.036 |
| *Primula auricula* | 1 | 0.003 | 4.061 | 0.010 |
| *Koeleria pyramidata* | 1 | 0.002 | 3.115 | 0.037 |
| *Veronica beccabunga* | 1 | 0.003 | 3.531 | 0.001 |
| *Omalotheca sylvaticum* | 1 | 0.002 | 3.307 | 0.001 |
| *Sagina saginoides* | 1 | 0.002 | 3.337 | 0.038 |
| *Pimpinella saxifraga* | 1 | 0.003 | 3.600 | 0.001 |
| *Cerastium holosteoides* | 1 | 0.003 | 3.546 | 0.032 |
| *Thymus praecox* subsp. *praecox* | 1 | 0.002 | 3.204 | 0.036 |
| Residual | 455 | 0.323 | | |

**Table S15** Metrics of niche divergence computed from 14 environmental predictors of the fine-grained dataset, including community-weighted means of six Karrer indicator values, for all pairwise comparisons among study species and between *Luzula alpina* (ALP) and the combined niche of *L. exspectata* (EXS) and *L. multiflora* (MUL). D: Schoener's *D*, I: Hellinger's *I*, $P_{Deq}$: *P*-value of the niche equivalency test for *D*, $P_{Ieq}$: *P*-value of the niche equivalency test for *I*, $P_{Dsim}$: *P*-value of the niche similarity test for *D*, $P_{Isim}$: *P*-value of the niche similarity test for *I*, $B_{PC1}$: niche breadth along PC1 (median ± standard deviation), $B_{PC2}$: niche breadth along PC2, S: niche stability, U: niche unfilling, E: niche expansion. Significant *P*-values < 0.05 are highlighted in bold. Niche breadth is given for the species indicated.

| **Comparison** | **D** | **I** | $P_{Deq}$ | $P_{Ieq}$ | $P_{Dsim}$ | $P_{Isim}$ | $B_{PC1}$ | $B_{PC2}$ | **S** | **U** | **E** |
|---|---|---|---|---|---|---|---|---|---|---|---|
| MUL – EXS | 0.71 | 0.88 | **0.0020** | **0.0010** | 0.3157 | 0.3377 | EXS 1.53±0.08 | EXS 1.23±0.10 | 0.97 | 0.15 | 0.03 |
| EXS – ALP | 0.67 | 0.89 | **0.0020** | **0.0020** | 0.1728 | 0.1728 | ALP 1.69±0.11 | ALP 1.24±0.11 | 0.91 | 0.00 | 0.09 |
| MUL – ALP | 0.75 | 0.93 | **0.0460** | **0.0180** | 0.1698 | 0.2368 | MUL 1.80±0.10 | MUL 1.40±0.10 | 0.96 | 0.03 | 0.04 |
| EXS+MUL – ALP | 0.74 | 0.94 | **0.0010** | **0.0060** | 0.0969 | 0.1279 | 1.74±0.10 | 1.38±0.11 | 0.98 | 0.02 | 0.02 |

**Table S16** Metrics of niche divergence computed from 14 environmental predictors of the fine-grained dataset, including community-weighted means of six Karrer indicator values, for all pairwise comparisons among study species and between *Luzula alpina* (ALP) and the combined niche of *L. exspectata* (EXS) and *L. multiflora* (MUL). D: Schoener's *D*, I: Hellinger's *I*, $P_{Deq}$: *P*-value of the niche equivalency test for *D*, $P_{Ieq}$: *P*-value of the niche equivalency test for *I*, $P_{Dsim}$: *P*-value of the niche similarity test for *D*, $P_{Isim}$: *P*-value of the niche similarity test for *I*, $B_{PC1}$: niche breadth along PC1 (median ± standard deviation), $B_{PC2}$: niche breadth along PC2, S: niche stability, U: niche unfilling, E: niche expansion. Significant *P*-values < 0.05 are highlighted in bold. Niche breadth is given for the species indicated.

| **RDA ANOVA model** | **Df** | **Variance** | **F value** | ***P*-value** |
|---|---|---|---|---|
| Model | 5 | 0.069 | 14.21 | 0.001 |
| Residual | 499 | 0.485 | | |
| **RDA ANOVA axes** | **Df** | **Variance** | **F value** | ***P*-value** |
| RDA1 | 1 | 0.052 | 53.969 | 0.001 |
| RDA2 | 1 | 0.017 | 17.504 | 0.001 |
| Residual | 502 | 0.485 | | |
| **RDA ANOVA explanatory variables** | **Df** | **Variance** | **F value** | ***P*-value** |
| Reaction (R) | 1 | 0.035 | 35.944 | 0.001 |
| Elevation | 1 | 0.017 | 17.969 | 0.001 |
| CryptogamCover | 1 | 0.009 | 9.173 | 0.001 |
| OrganicCover | 1 | 0.004 | 4.401 | 0.011 |
| Northness | 1 | 0.003 | 3.559 | 0.036 |
| Residual | 499 | 0.486 | | |

**Methods S1** Isolation-by-distance and isolation-by-environment

We explored the impact of demographic and ecological processes on the spatial structure of genetic variation by computing isolation-by-distance (IBD) and isolation-by-environment (IBE) for each species (Wang *et al.*, 2013). Following Huynh *et al.* (2020), we inferred patterns of IBD by comparing Euclidean geographic and genetic distances (rho/(rho-1) with Mantel tests using 20,000 permutations in *vegan*. Signatures of IBE are expected if gene flow occurs predominantly among individuals from similar ecological conditions (Wang & Bradburd, 2014) and were tested through partial Mantel tests of genetic distances and Euclidean distances from principal components (PCs) of environmental space derived from the coarse-grained dataset while accounting for spatial structure. We validated patterns of IBD and IBE through partial RDA with 20,000 permutations, associating population-level allele frequencies with coordinates along the first two PCs of environmental space while accounting for geographical isolation.

**Methods S2** Environmental data acquisition

We combined climatic, topographic, lithologic and edaphic data to assess differences in coarse-grained abiotic ecological niches among our three study species. Climatic variables were retrieved at 30-arc second resolution from CHELSA (Karger *et al.*, 2017). A digital elevation model (DEM) for Europe at 30 m resolution was obtained from the GMES RDA project (EU-DEM; http://data.europa.eu/88u/dataset/f576cda8-d598-478c-b8fe-ad2634c927e8) and used to compute the topographic variables elevation, slope, northness and topographic roughness, which were aggregated to 1 km resolution. Lithological data from Geo-LiM (Donnini *et al.*, 2020) were used to construct a bedrock-type layer following Chauvier *et al.* (2021). First, soil lithology was classified into three categories: calcareous, siliceous, and mixed (Table Bedrock Classes). In a second step, this newly created layer was converted to a 100 m grid for each category. Finally, all 100 m grids were aggregated to 1 km resolution by averaging the proportions of the three categories within each grid cell. Topsoil data that are ecologically relevant for plant species were retrieved from SoilGrids (Hengl *et al.*, 2017). Climatic data for the Last Glacial Maximum used to project ENMs to paleoclimatic conditions were retrieved from CHELSA for four general circulation models (IPSL-CM5A-LR, CCSM4, MRI-CGCM3, MPI-ESM-P).

**Bedrock Classes**: Conversion of Geo-LIM lithological categories into bedrock classes and corresponding bedrock acidity (ranging from 0 to 1).

| Geo-LIM category | Bedrock class | Bedrock acidity |
|---|---|---|
| acid rocks | siliceous | 1 |
| claystone | siliceous | 1 |
| mafic rocks | mixed | 0.5 |
| sandstone | siliceous | 1 |
| intermediate rock | mixed | 0.5 |
| gypsum evaporite | calcareous | 0 |
| mix carbonate rocks | calcareous | 0 |
| pure carbonate rocks | calcareous | 0 |
| metamorphic rocks | mixed | 0.5 |

**Results S1** Analyses of vegetation data

The communities of accompanying plant species were strongly overlapping among our study species, as reflected by the lack of separation in the DCA and TWINSPAN clusters that did not align with the three *Luzula* species (Fig. 4a). Characteristic species for the two major TWINSPAN clusters were *Nardus stricta* on the one hand, and *Bistorta vivipara*, *Carex sempervirens*, *Galium anisophyllon*, *Lotus corniculatus*, *Sesleria caerulea* and *Trifolium pratense* on the other hand. The first and larger (n = 356) TWINSPAN cluster consisted mainly of ALP (65%), whereas MUL (19%) and EXS were scarcer (15%). Even though ALP was still the dominant species (49%) of the second and smaller (n = 149) cluster, EXS reached a comparable prevalence (42%), while MUL remained rare (9%). Most individuals of ALP (76%) and MUL (83%) belonged to the first TWINSPAN cluster, whereas accessions of EXS were more evenly distributed between the two clusters (47% and 53%, respectively.)

Although the accompanying plant communities were overall similar among the three study species, redundancy analysis (RDA) revealed a number of diagnostic species, which in case of

EXS were mostly characteristic for alpine grasslands over limestone (e.g., *Betonica alopecurus*, *Koeleria pyramidata*, *Potentilla crantzii*). In contrast, occurrences of ALP were characterised by the presence of species typical for acidic alpine meadows (e.g., *Helictochloa versicolor*, *Nardus stricta*, *Potentilla erecta*), and MUL co-occurred with acidophilic species of alpine dwarf-shrub heaths (e.g., *Kalmia procumbens*, *Vaccinium myrtillus*, *Vaccinium vitis-idaea*).

Together, the environmental preferences of our study species inferred from both coarse- and fine-grained analyses were consistent with previous findings (Bačič *et al.*, 2019; Geurden *et al.*, 2025) and characterised EXS as a species of alpine grasslands on calcareous substrates in areas with high snowfall. In contrast, MUL thrives in dwarf-shrub heaths with more acidic soil and better nitrogen availability. Lastly, ALP occurs predominantly in alpine meadows at high elevations over siliceous bedrock.

**Results S2** Idiosyncratic patterns of refugial areas and range expansion

Combining evidence from species distribution modelling, spatial patterns of $\psi$ and private alleles, we inferred mostly peripheral refugia over limestone for EXS (Figs. 2, 3), which is in line with its preference for calcareous bedrock. These refugia were distributed partly along the southeastern margin of the Alps, a region that acted as an important Pleistocene refugium for many plant species (Tribsch & Schönswetter, 2003; Schönswetter *et al.*, 2005). In contrast, suitable habitat during the LGM coincided with known refugia for silicicolous plants in the Eastern Alps in case of MUL, but was more widespread along the periphery of the Alps for ALP, and spatial patterns of allele frequencies supported these results. Recolonisation of the Alps from glacial refugia followed idiosyncratic patterns for the three species as inferred from spatial patterns of $\psi$ and $\pi$ (Fig. 2). EXS expanded from the southern margin of the Alps, whereas the spread of ALP likely started somewhere along the western margin of the Eastern Alps, suggesting that the species may have persisted also within refugia in the Western Alps. Finally, MUL did not expand far beyond its refugial areas. All three species exhibited low genetic differentiation among populations which is likely explained by wind-pollination enabling gene flow across large distances (Kuzmanović *et al.*, 2017).
